# Multiscale analysis of acne connects molecular subnetworks with disease status

**DOI:** 10.1101/587857

**Authors:** Jacob B. Hall, Aparna A. Divaraniya, Hao-Chih Lee, Christine E. Becker, Benjamin McCauley, Patricia K. Glowe, Robert Sebra, Ana B. Pavel, Giselle Singer, Amanda Nelson, Diane Thiboutot, Ellen Marmur, Eric E. Schadt, Joshua Zeichner, Emma Guttman-Yassky, Brian A. Kidd, Joel T. Dudley

## Abstract

Acne vulgaris affects millions of individuals and can lead to psychosocial impairment as well as permanent scarring. Previous studies investigating acne pathogenesis have either examined a targeted set of biological parameters in a modest-sized cohort or carried out high-throughput assays on a small number of samples. To develop a more comprehensive understanding of acne pathophysiology, we conducted an in-depth multi-omic study of 56 acne patients and 20 individuals without acne. We collected whole blood, skin punch biopsies, microbiota from skin follicles, and relevant clinical measurements to understand how multiple factors contribute to acne. We provide an integrative analysis of multi-omics data that results in a molecular network of acne. Comparisons of lesional and non-lesional skin highlighted multiple biological processes, including immune cell and inflammatory responses, response to stress, T cell activation, lipid biosynthesis, fatty acid metabolism, keratinocytes, antimicrobial activity, epithelial cell differentiation, and response to wounding, that are differentially altered in acne lesions compared to non-lesions. Our results suggest baseline differences in the skin that may predispose individuals to develop acne. These datasets and findings offer a framework for new target identification and reference for future studies.

## INTRODUCTION

Acne vulgaris is a common inflammatory skin disease that affects approximately 39 million people in the United States and about 615 million people globally (Vos *et al*., 2017). While primarily afflicting teenagers, over 54% of adults are also affected beyond their teenage years (Institute for Health Metrics and Evaluation (IHME); Lim *et al*., 2017). The episodic, and sometimes chronic, nature of disease results in considerable economic, medical, and psychological burdens (Behnam *et al*., 2013; Lim *et al*., 2017; Picardi *et al*., 2013). Addressing this complex condition warrants a more comprehensive picture of disease from multiple dimensions of data.

Acne develops from a combination of four critical factors: (i) increased sebum production, (ii) abnormal keratinization and cornification of the sebaceous follicular duct, (iii) colonization of the hair follicles by microbes—putatively *Cutibacterium acnes (C. acnes;* formerly *Propionibacterium acnes*, or *P. acnes*), and (iv) a localized inflammatory response to various host and environmental mediators, including free fatty acids produced by *C. acnes* and sensitized immune cells (Lichtenberger *et al*., 2017; Tuchayi *et al*., 2015; Zouboulis *et al*., 2005). Despite determining factors that drive pathogenesis, the precise details of their molecular and cellular relationships remain unclear, particularly in how these components shape the immune system and modulate the immune response in acne lesions and healthy skin (Dreno *et al*., 2015; Mattii *et al*., 2018; Squaiella-Baptistão *et al*., 2015). Elucidating how lipid production, keratinization, inflammatory mediators, and microbes contribute to acne lesions could uncover disease subtypes and suggest new treatment strategies.

We conducted a comprehensive multi-omics study of acne vulgaris to investigate the molecular and cellular factors associated with inflammation in patients with and without active acne. To evaluate local (skin) and systemic (blood) mediators, we examined high-throughput measurements from multiple biological levels, including the genome, transcriptome, cytome, and microbiome, for 56 individuals with active mild-to-moderate acne and 20 age-matched controls without acne. We applied integrative analysis and constructed the first molecular network of acne lesions. We highlight differentially expressed genes and co-expressed genes (“modules”) that appear to drive inflammation in acne lesions, as well as genes that display differential expression in non-lesional skin in acne patients compared to controls. These results describe new factors that affect acne pathogenesis and point to possible new therapeutic targets worthy of further investigation.

## RESULTS

### Subject characteristics and study overview

We enrolled 56 adults with mild-to-moderate acne, and 20 age- and sex-matched non-acne controls (**Table 1**, enrollment criteria described in **Text S1**). Participants included multiple racial backgrounds and skin types. We observed no associations with disease and demographic factors, including age, sex, ethnicity, and Fitzpatrick skin score (**Table 1**; **Table S1**).

**Table 1.**
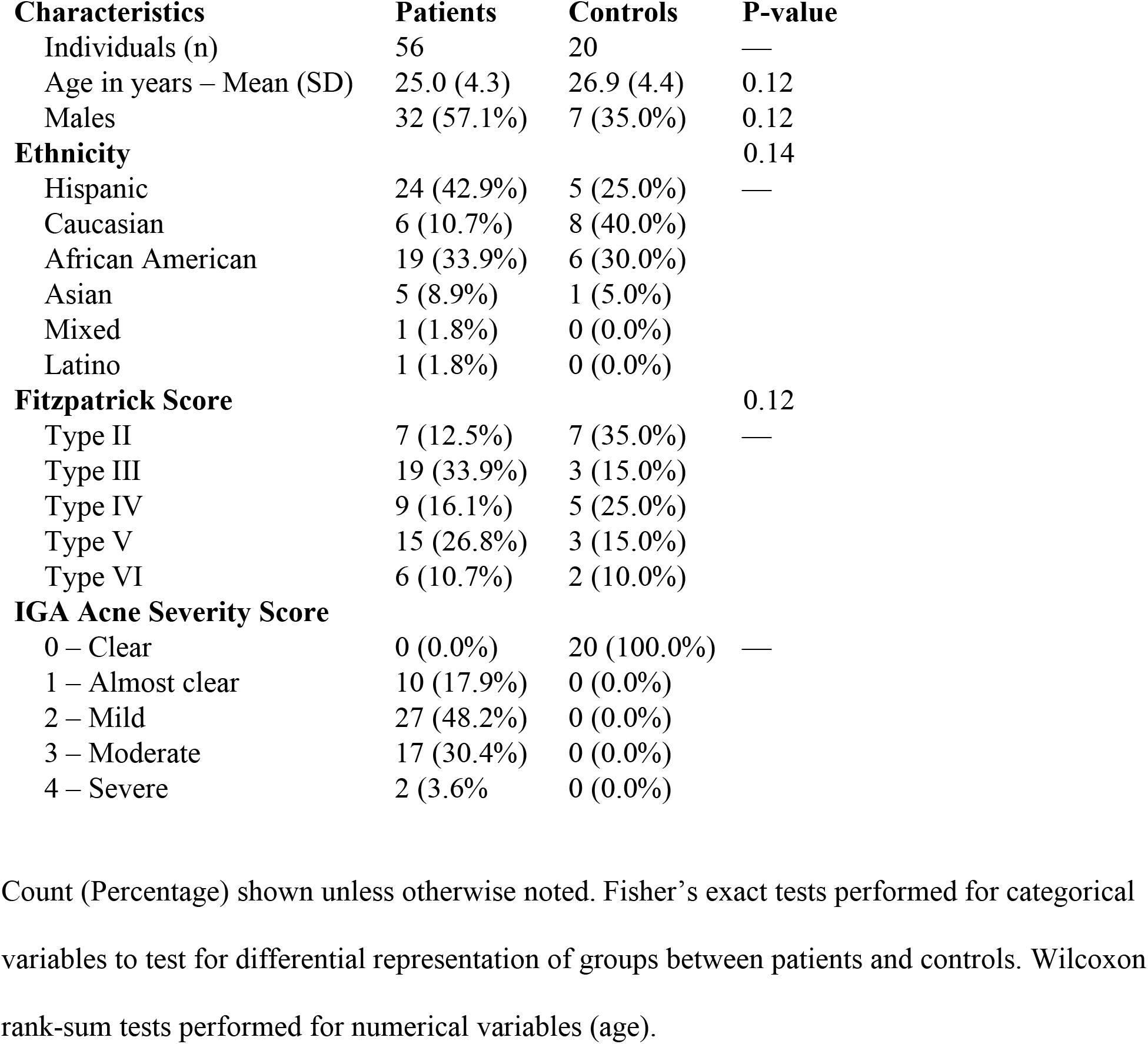
Cohort characteristics.

| **Characteristics** | **Patients** | **Controls** | **P-value** |
| --- | --- | --- | --- |
| Individuals (n) | 56 | 20 | — |
| Age in years – Mean (SD) | 25.0 (4.3) | 26.9 (4.4) | 0.12 |
| Males | 32 (57.1%) | 7 (35.0%) | 0.12 |
| **Ethnicity** |  |  | 0.14 |
| Hispanic | 24 (42.9%) | 5 (25.0%) | — |
| Caucasian | 6 (10.7%) | 8 (40.0%) |  |
| African American | 19 (33.9%) | 6 (30.0%) |  |
| Asian | 5 (8.9%) | 1 (5.0%) |  |
| Mixed | 1 (1.8%) | 0 (0.0%) |  |
| Latino | 1 (1.8%) | 0 (0.0%) |  |
| **Fitzpatrick Score** |  |  | 0.12 |
| Type II | 7 (12.5%) | 7 (35.0%) | — |
| Type III | 19 (33.9%) | 3 (15.0%) |  |
| Type IV | 9 (16.1%) | 5 (25.0%) |  |
| Type V | 15 (26.8%) | 3 (15.0%) |  |
| Type VI | 6 (10.7%) | 2 (10.0%) |  |
| **IGA Acne Severity Score** |  |  |  |
| 0 – Clear | 0 (0.0%) | 20 (100.0%) | — |
| 1 – Almost clear | 10 (17.9%) | 0 (0.0%) |  |
| 2 – Mild | 27 (48.2%) | 0 (0.0%) |  |
| 3 – Moderate | 17 (30.4%) | 0 (0.0%) |  |
| 4 – Severe | 2 (3.6%) | 0 (0.0%) |  |

Count (Percentage) shown unless otherwise noted. Fisher’s exact tests performed for categorical variables to test for differential representation of groups between patients and controls. Wilcoxon rank-sum tests performed for numerical variables (age).

For each subject, we collected skin biopsies from the upper back, follicular contents from the facial cheek, and peripheral blood to generate a comprehensive biological profile of local skin and non-local, peripheral blood (**Figure 1**). We obtained 4-mm full-thickness skin biopsies from both active acne lesional sites and proximal non-lesional sites (collection described in **Text S2**, examples shown in **Figure S1**). For controls without active acne, we collected a single 4-mm skin biopsy from a similar location as the acne patients’ biopsies. We obtained micro-comedone samples via cyanoacrylate glue follicular biopsies to assess the skin microbiome (see **Materials and Methods; Text S2**).

**Figure 1.**
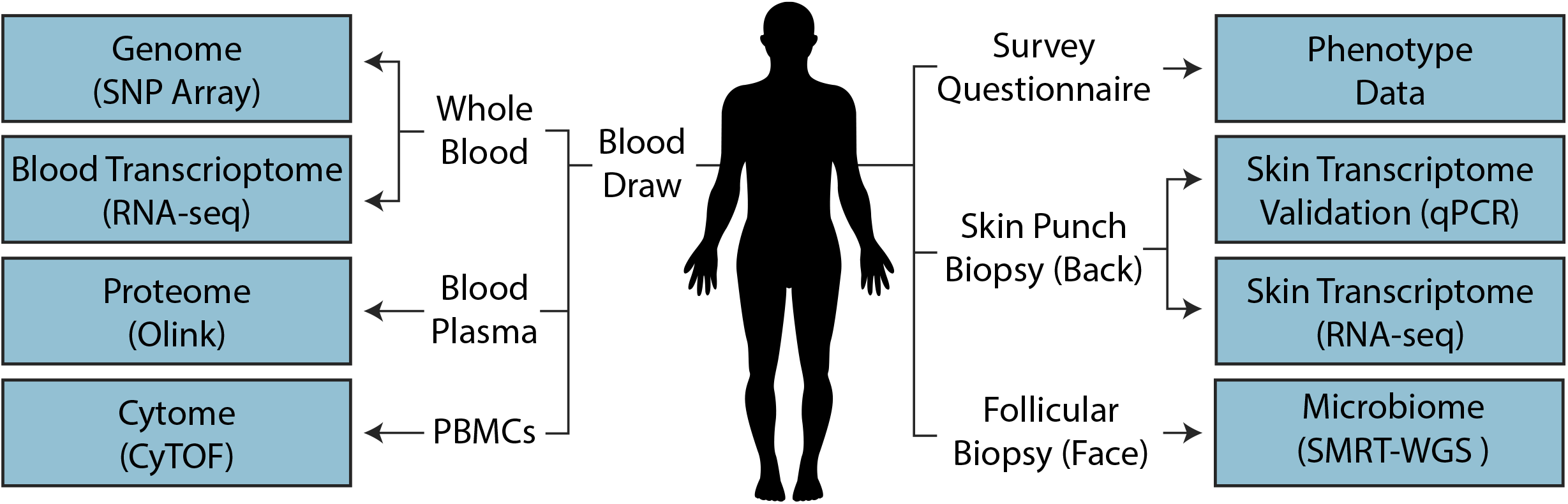
Study design. Data types collected and corresponding assays performed. PBMCs, Primary Blood Monocyte Cells; qRT-PCR, quantitative polymerase chain reaction; RNA-seq, RNA sequencing; SNP, Single Nucleotide Polymorphism; CyTOF, Cytometry by Time of Flight.

### Peripheral blood does not provide an acne-specific disease signature

To determine whether a biomarker signature of acne existed in peripheral blood, we collected whole blood samples from each person and measured single nucleotide polymorphisms (SNPs), proportions of specific cell types, levels of plasma proteins, and gene transcript counts from whole-genome RNA sequencing (RNA-seq) (**Figure 1**). We interrogated over 1 million genotyped SNPs and none exhibited genome-wide significance between acne patients and controls (**Figure S2**). A targeted subset of 306 SNPs within ± 30 kilobases of the 27 loci associated with acne vulgaris in the GWAS catalog (MacArthur *et al*., 2017) (accessed July 2017) also showed no genotype variants that were significantly associated with acne case status. We examined thousands of single-cells using time of flight mass cytometry (CyTOF), yet found no significant differences in the frequencies of twelve immune cell subsets between acne patients and controls (**Figure S3**). Comparison of gene expression levels from RNA-seq of whole-blood cells found no genes with statistical differences in expression between case status (**File S1a**). Lastly, a panel of 92 immune response and inflammatory proteins in blood plasma exhibited no significant differences in abundance between patients and controls (**Table S2**).

### Keratinocyte proliferation genes exhibit baseline differences between patients and controls

To identify molecular differences in unaffected skin tissue, we analyzed RNA expression in non-lesional skin biopsy samples from patients and controls. We profiled more than 17,500 unique transcripts and found 68 genes differentially expressed (FDR < 5%) (**Figure 2a; File S1a**). Functional enrichment of genes over-expressed in non-lesional skin from patients compared to controls highlighted pathways related to keratinization and immune response, including keratin genes involved in tissue development (*KRT6A, KRT16*), a transcription factor that regulates epithelial cell differentiation (*EHF*), genes with calcium- and zinc-binding motifs that mediate the inflammatory response to wounding (*S100A8, S100A9*), and a superoxide dismutase (*SOD2*) associated with protein homotetramerization to detoxify reactive oxygen species produced during immune response (Fisher’s exact test, p < 0.05, FC > 13) (**File S1b; Figure 3b**).

**Figure 2.**
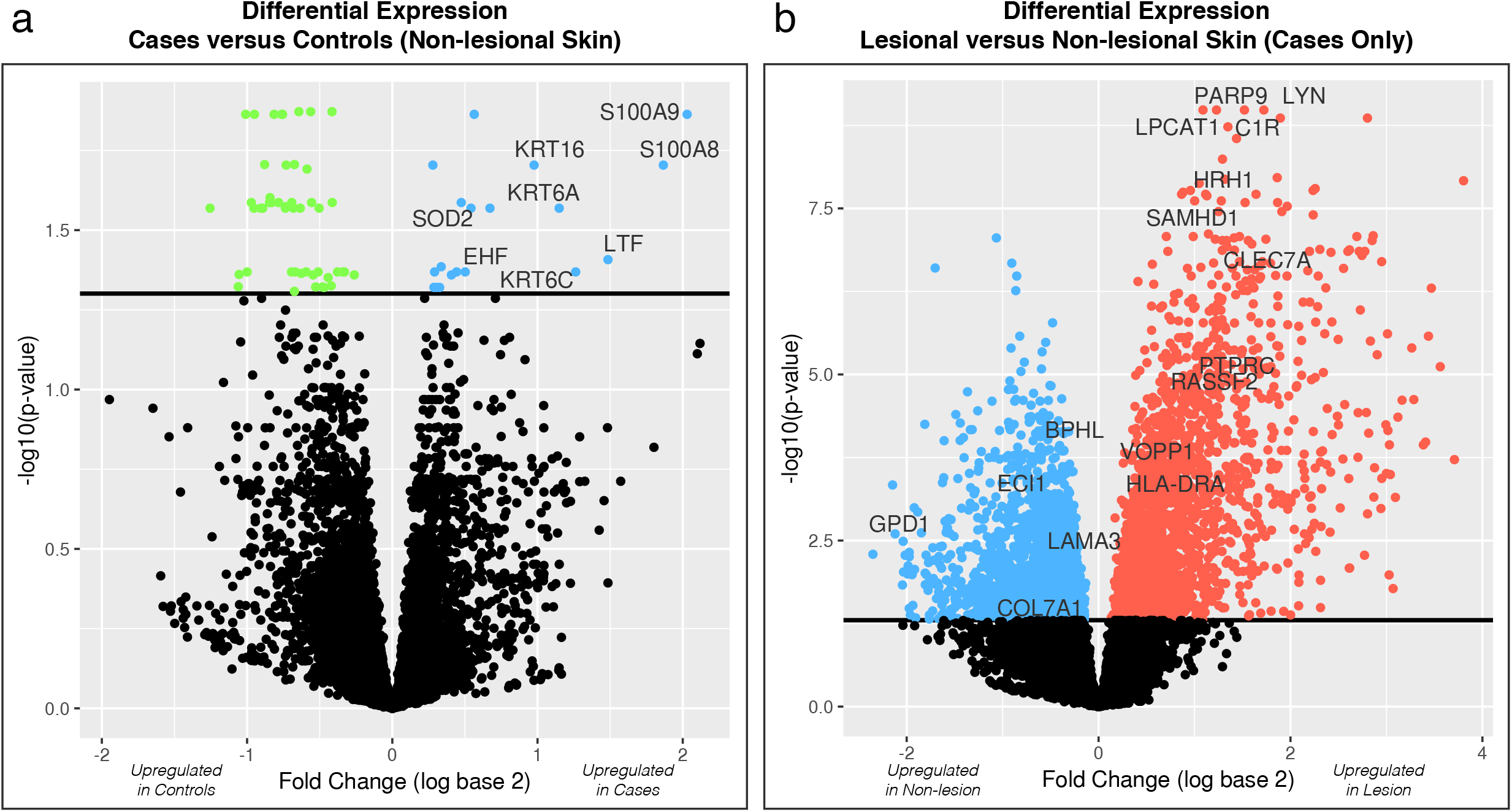
Differentially expressed genes. Volcano plots for **(a)** case/control differential expression comparisons for 17,598 genes and **(b)** lesion/non-lesion differential expression comparisons for 17,713 genes. Solid horizontal lines represent significance threshold for FDR < 5%. Each point represents a single gene. Points above the significance threshold are colored based on upregulation, as labeled on the x-axis. Select genes described throughout the study are annotated.

**Figure 3.**
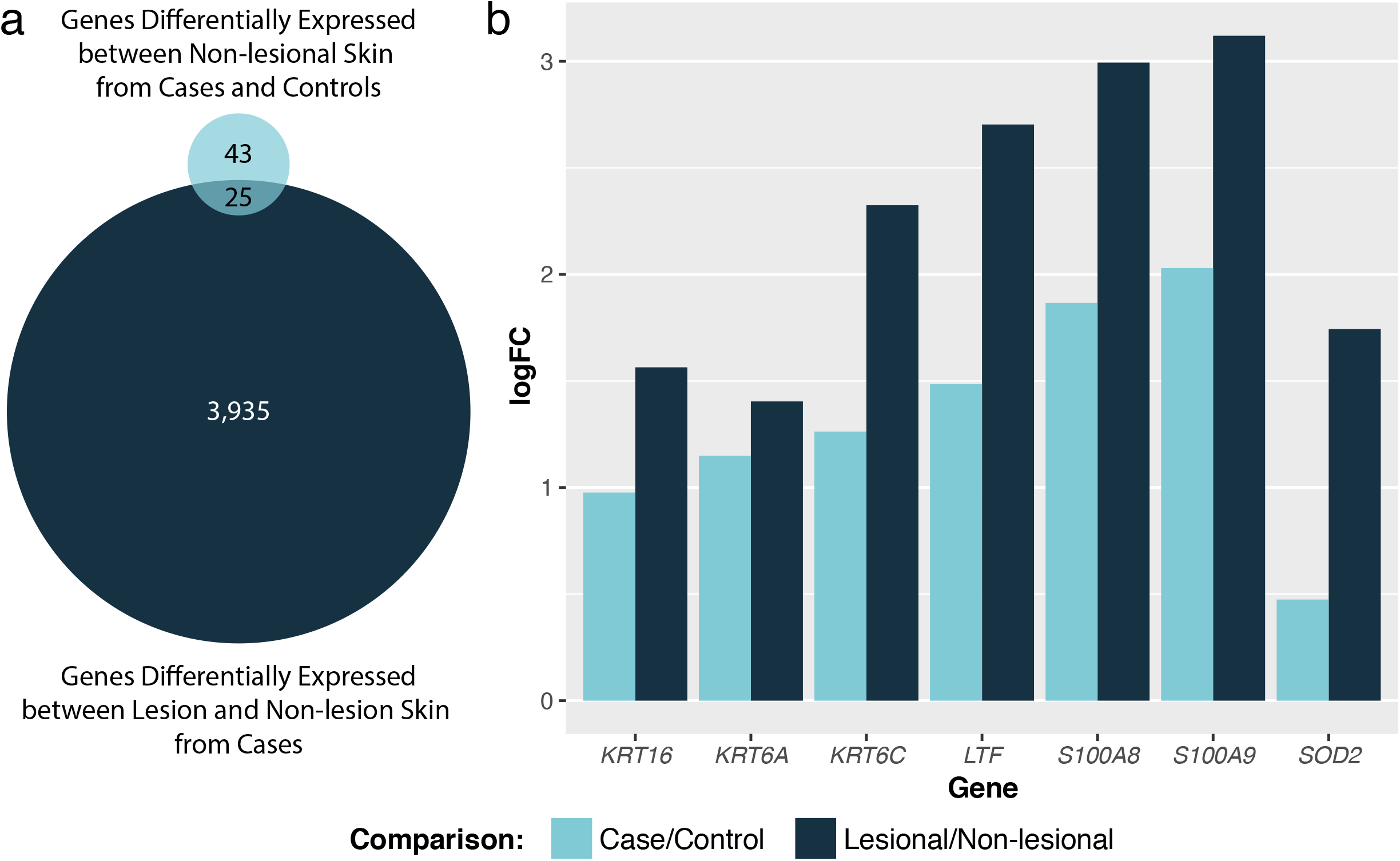
Differentially expressed genes upregulated in lesion samples and in patients compared to controls. **(a)** Venn diagram of significant differentially expressed genes for lesion/non-lesion and case/control comparisons. **(b)** Paired bar plots for seven select genes that are differentially expressed in both analyses. Greater fold changes are observed for the lesion/non-lesion comparison for each gene. Log fold change (logFC) were calculated from expression measured using RNA-seq.

### Immune response and inflammatory mediator genes are differentially expressed between lesional and non-lesional skin from acne patients

To contrast the molecular profiles between active acne lesion sites and non-lesional skin, we characterized genome-wide RNA expression measurements of paired biopsy samples from acne patients. We examined more than 17,500 unique transcripts and found 3,960 differentially expressed genes (FDR < 5%), with 2,286 genes upregulated in lesional compared to non-lesional skin (**Figure 2b; File S1a**). Twenty-five of these differentially expressed genes were also differentially expressed in the case-control comparison (**Figure 3a, Table S3**). Rank-ordering the upregulated genes by p-value highlights genes that regulate innate and adaptive immune responses (*LYN*), initiate the classical complement cascade (C1R), help repair DNA damage and modulate an immune response through interferon-mediated antiviral defenses (*PARP9*), and metabolize phospholipids (*LPCAT1*) (**Figure 2b; File S1a**).

To better understand the transcriptional changes associated with active disease, we applied functional enrichment analysis to differential expression profiles (see **Materials and Methods**). Genes over-expressed in acne lesions were enriched for defense response (e.g. *SAMHD1*, Triphosphohydroslase 1, which promotes TNF-alpha protoinflammatory response), immune response (e.g. *IL4R*, Interleukin 4 Receptor, which promotes Th2 differentiation), inflammatory response (e.g. *CCR1*, C-C Motif Chemokine Receptor 1, which is a receptor for macrophage inflammatory protein 1 alpha), response to stress (e.g. *BCL3*, B cell CLL / Lymphoma 3, which activates transcription through association with NF-kappa B), leukocyte activation (e.g. *CLEC7A*, which binds and promotes proliferation of T cells in a way that does not involve their surface glycans), regulation of cytokine production (e.g. *TLR8*, which stimulates cytokine secretion), and T cell activation (e.g. *CD7*, which is found on mature T cells and plays an essential role in T cell interactions) (Fisher’s exact test, p < 3 × 10^−9^, FC > 2.2) (**File S1b**). Genes under-expressed in acne lesions were enriched for lipid metabolism and biosynthesis, steroid metabolism and biosynthesis, and fatty acid metabolism (Fisher’s exact test, p < 4 × 10^−6^, FC > 2.5) (**File S1b**).

### Co-expressed genes influencing immune response and lipid metabolism linked with active acne

To evaluate correlated patterns of gene expression in active acne, we applied weighted gene coexpression network analysis (WGCNA) to construct a molecular network of gene expression measured from lesional skin samples. We used a soft-threshold R^2^ > 0.80 to determine genes with correlated expression and found 49 unique sets of genes (**File S1f**), defined as “modules”, that were co-expressed. Of these 49 modules, 21 showed significant overlap (OR > 1, FDR p-value < 5%) with upregulated (in lesion or non-lesion) differentially expressed gene sets (**File S1c**). We conducted enrichment analysis to understand the functional relationships of the genes within each module and found modules for immune and defense response, fatty acid oxidation, cell cycle regulation, cell signaling, lipid metabolism, and lipid biosynthesis (**File S1d,e,g**).

Some of the most significantly differentially expressed genes for the fatty acid metabolism module, lesion module 05 (LM05), include genes such as *ECI1* (Enoyl-CoA Delta Isomerase 1), which is involved in beta-oxidation of unsaturated fatty acids, *GPD1* (Glycerol-3-Phosphate Dehydrogenase 1), which plays a critical role in carbohydrate and lipid acid metabolism, and *BPHL* (Biphenyl Hydrolase Like), which may play a role in cell detoxification. Significant genes from the defense response module (LM01) include genes like *SAMHD1* (Triphosphyohydrolase 1), which may be involved in tumor necrosis factor-alpha proinflammatory response, and *CLEC7A* (C-Type Lectin Domain Containing 7A), which stimulates T cell proliferation.

A closer look at the T cell receptor signaling and cell communication module, LM10 (**Figure 4a**), identified four hub genes (highly co-expressed) — *RASSF2* (Ras association domain family member 2), *HLA-DRA* (major histocompatibility complex class II DR alpha), *VOPP1* (vesicular overexpressed in cancer prosurvial protein 1), and *PTPRC* (protein tyrosine phosphatase receptor type C) — that were also differentially expressed in lesional versus non-lesional skin samples (**File S1f**). *HLA-DRA* and *PTPRC* control immune cell response through antigen presentation and T cell activation, respectively.

**Figure 4.**
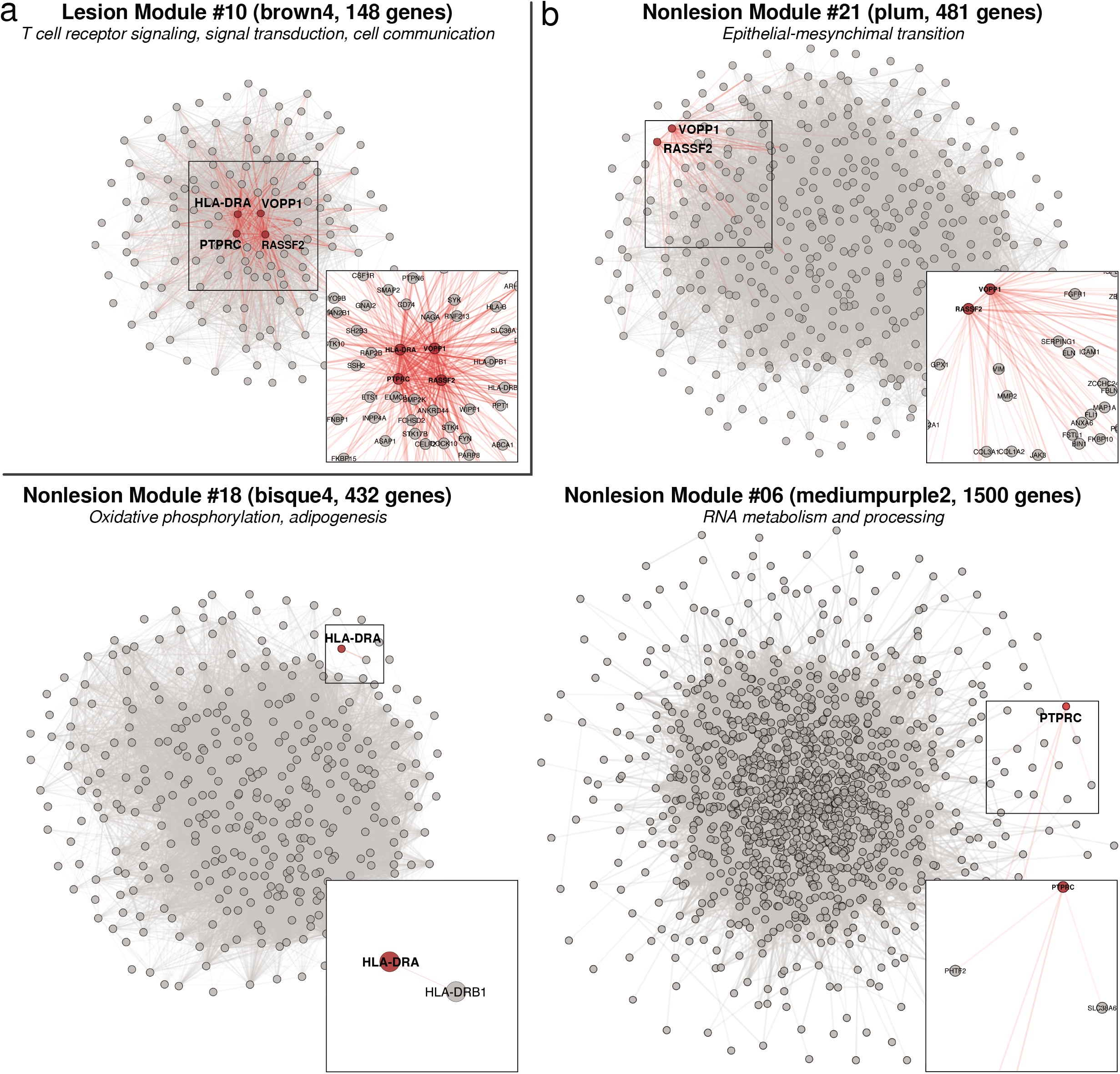
Co-expression module connectivity comparison. Comparison of intramodule connectivity for modules containing *HLA-DRA, RASSF2, PTPRC*, and *VOPP1* in the **(a)** lesion and **(b)** multiple non-lesion co-expression networks. Circles/nodes represent genes and lines/edges represent co-expression between genes. The four highlighted genes are shown using red circles. All gene-gene connections with correlation R^2^ > 0.1 are shown. Red lines represent all connections involving the four highlighted genes. Grey lines represent all other connections. Line transparency of all lines is scaled to the strength of the gene-gene connections. Subset panels show zoomed-in portions of modules to highlight genes that are co-expressed with the highlighted genes; uninvolved genes and connections are hidden in subset panels. Created in Cytoscape v3.6.1, with node layout set to ‘perfuse force directed’.

### Hub genes in lesion skin network are co-expressed differently in non-lesional skin

To explore differences in the molecular networks of inflamed and un-inflamed skin in acne patients, we also generated a co-expression network of genes expressed in non-lesional skin. We observed 52 unique modules (**File S1f**), 16 of which showed significant overlap of differentially expressed genes (OR > 1, FDR p-value < 5%) with those observed in lesional skin, compared to non-lesional skin (**File S1c**). Functional enrichment of these modules implicated pathways such as epidermis development, immune response, cell cycle regulation, cell adhesion, extracellular matrix organization, and ion/cation transport (**File S1d, e, g**). Top significant differentially expressed genes from the epidermis development module non-lesion module 04 (NLM04) include *LAMA3*, which is essential for formation and function of the basement membrane and helps regulate cell migration and mechanical signal transduction, and *COL7A1*, which functions as an anchoring fibril between the external epithelia and the underlying stroma.

The four hub genes in LM10 altered their intra-module connectivity completely in the non-lesional skin tissue, especially for *HLA-DRA* and *PTPRC* (**Figure 4b**). We observed lower correlation between these gene’s expression values and found them in separate modules in the non-lesion skin network (**Figure 4b**), suggesting *RASSF2, VOPP1, HLA-DRA*, and *PTPRC* play a greater role in modulating T cell receptor signaling and immune responses in lesional compared to non-lesional skin. The most similar (“preserved”) module in the non-lesion network, compared to genes making up LM10, was NLM37 (black) (**File S1h**), which is enriched for genes that regulation transcription (**File S1g**).

### Skin-associated modules highlight genes affecting epidermal differentiation

We examined the expression patterns of skin-associated modules to better understand the biological processes involved in acne pathogenesis. We found five lesion skin modules enriched for genes associated with skin-related processes, including keratinocyte differentiation and epidermis development (LM11, LM17, LM47), skin epidermal cells (LM22), and melanogenesis (LM18) (**File S1g**). One keratinocyte differentiation module (LM11) included genes for calcium binding and small proline rich proteins overexpressed in lesion samples (*S100A7, SPRR2A, SPRR2B, SPRR2F*, and *SPRR2G*). The skin epidermal cell module (LM22) contained 24 genes associated with CD200+CD49+ hair follicle keratinocytes (McCall *et al*., 2014) that were overexpressed in the non-lesional skin from acne patients (Fisher’s exact test, p < 1 × 10^−6^, FC > 3.5) (**File S1e**). Lastly, LM17 included keratin and collagen genes associated with epidermis development that were overexpressed in the non-lesional skin of patients (Fisher’s exact test, p < 1.2 × 10^−5^, FC > 7.8) (**File S1e**). These results suggest that differences in expression levels of genes that promote keratinization and epidermal development are already present at the non-lesional skin of acne patients.

### Independent validation of gene expression using qRT-PCR

To validate expression differences observed from RNA-seq, we selected several of the top differentially expressed genes to measure independently using qRT-PCR. We chose the four hub genes described from LM10, as well as genes that play roles in innate/anti-bacterial immune responses, T cell activation, antigen-presentation, lipid metabolism, and keratinocyte differentiation and proliferation; these included: *HLA-DRA, HK3, C1R, LPCAT1, LYN, PARP9, PTPRC, RASSF2, and VOPP1*. All of these genes showed significant increases in lesional acne skin as compared with both normal (skin from controls) and non-lesional acne skin (**Figure S4**). We found significant concordance between the RNA-seq and PCR results (Fisher’s exact test, p < 0.02) (**Table S4**).

### *Cutibacterium acnes* is more abundant in patients with acne

To assess potential differences in the facial microbiomes of acne and control subjects, we conducted metagenomic sequencing of DNA isolated from cyanoacrylate glue follicular biopsies (Hall *et al*., 2018). We used long-read single-molecule real time (SMRT) RSII sequencing technology from Pacific Biosciences (PacBio) for microbiome samples from acne patients, non-acne controls, a positive mock community control, and a negative sampling method control. After quality control steps to ensure sufficient abundance of high-quality microbial sequence reads, 20 acne patients and 7 controls remained for metagenomic analysis (see **Materials and Methods; Figures S5–7**). We found distinct microbiome profiles associated with acne and control tissues (**Figure 5; Table S5**). Acne patients had significantly higher relative abundance of *C. acnes* than controls (58.9% vs 2.4%) (p = 0.0203) (**Table S5**). *Cutibacterium phage (C. phage*) were assessed cumulatively and had higher (though not statistically significant) mean relative abundance in patients (17.2%) compared to controls (7.0%) (p = 0.3043) (**Table S5**). Additionally, we attempted to determine specific ribotypes of *C. acnes* but were unable to due to a lack of sufficient sequence coverage of the *C. acnes* genome for the majority of samples.

**Figure 5.**
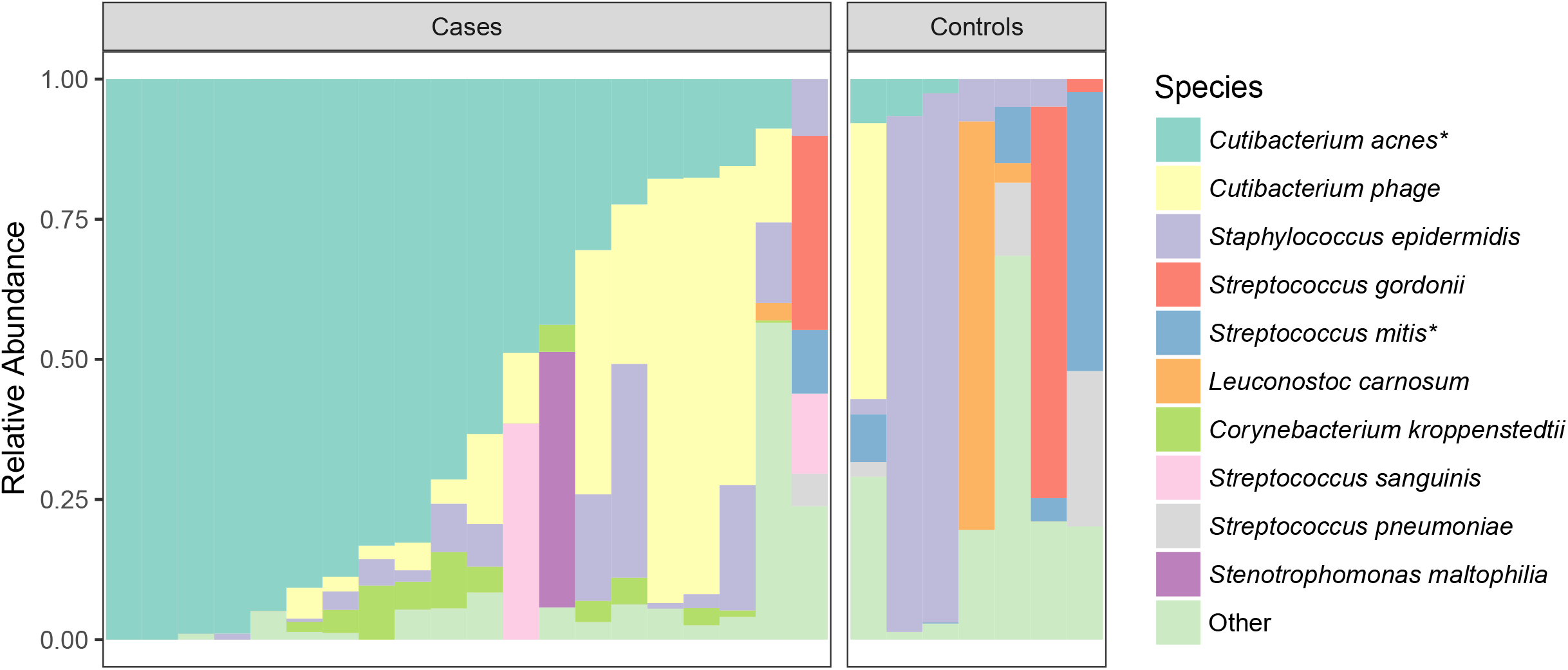
Microbiome profile of top species for acne patients and controls. Stacked bar chart showing relative species abundance for each sample (column), grouped by case status. The top ten species with the highest mean relative abundance cohort-wide are shown. All other species are grouped into the “Other” category. Samples are ordered left to right based on species abundance. Asterisks mark species with significant differential abundance between case and controls, controlling for multiple hypothesis testing (Wilcoxon text, adjusted p-value < 0.05).

## DISCUSSION

In this study, we evaluated potential molecular and cellular factors associated with acne by examining a collection of multi-omic data from 56 adults with active, mild-to-moderate acne, as well as 20 controls without acne. Overall, we found no blood biomarkers with significant differences between patients and controls, suggesting that acne is not a systemic condition, particularly for mild-to-moderate acne. In contrast, we found multiple molecular differences between active acne lesions and unaffected non-lesional skin for inflammatory responses, innate and antibacterial responses, lipid metabolism, and keratinization.

Keratinization plays an essential role in acne pathogenesis, affecting the accumulation of sebum and formation of comedones (Tanghetti, 2013). We found that three keratin genes associated with epidermal proliferation (*KRT6A, KRT6C*, and *KRT16*) were overexpressed in lesional skin of acne patients, compared to non-lesional skin. Epidermal proliferation has been shown to be altered due to activated inflammatory pathways in atopic dermatitis (Khattri *et al*., 2014). We also discovered differences in inflammatory and antimicrobial genes, which collectively suggest baseline differences in expression that may make skin cells more susceptible to abnormal keratinization and immune responses to microbes.

We generated co-expression networks for lesion and non-lesion skin biopsies from acne patients and annotated the biological processes for the gene sets comprising each module using functional enrichment analysis. Immune and inflammatory processes were linked to upregulated genes in lesional skin. In particular, we found four hub genes associated with lesional skin (*RASSF2, HLA-DRA, VOPP1*, and *PTPRC*). We hypothesize that targeting and modifying expression of these genes could alter the biological functions associated with this module, to approach a more non-lesional state. Using qRT-PCR, we confirmed abundant expression of all four genes and validated they were over-expressed in lesion skin compared to non-lesion skin in acne patients.

Two of the hub genes in LM10, *RASSF2* and *VOPP1*, regulate apoptosis. Maintaining control of programmed cell death is critical for preserving the balance between normal and abnormal skin remodeling of keratinocytes and sebocytes in response to environmental triggers, inflammation, and treatments, including isotretinoin (Melnik, 2017; Mrass *et al*., 2004; Nelson *et al*., 2008). The association between *RASSF2* and *VOPP1* expression and acne, and their role in apoptosis, points to new therapeutic targets worth investigating. Isotretinoin, the most effective treatment for severe acne, triggers apoptosis in sebaceous glands (Nelson *et al*., 2008). Elevated levels of *RASSF2* have been shown to increase apoptosis while higher levels of *VOPP1* may decrease apoptotic potential (Baras *et al*., 2011; Kanwal *et al*., 2017). We observed both genes were upregulated in acne lesion samples. Previous work has shown that isotrention therapy up-regulates *RASSF2* (Nelson *et al*., 2009), which suggests the precise mechanism of action, as well as the role that *RASSF2* may play, warrants further investigation for establishing better control of inflammatory mechanisms and apoptotic triggers without the teratogenicity and other possible side effects associated with isotretinoin use.

Acne pathogenesis arises from inappropriate responses of both the innate and adaptive immune systems (Koreck *et al*., 2003). Two additional hub genes in LM10, *HLA-DRA* and *PTPRC*, were over-expressed in acne lesions and may help regulate immune response. *HLA-DRA* codes for a protein on the surface of antigen presenting cells, including dendritic cells (Brooks and Moore, 1988). *PTPRC* controls T cell receptor signaling through dephosphorylation of *LCK* and *FYN* (Porcu *et al*., 2012; Stanford *et al*., 2012). Both *HLA-DRA* and *PTPRC* may serve as targets to regulate dendritic and T cells and promote a non-lesion skin expression profile.

We found preliminary evidence of a link between *C. acnes* abundance and the expression of differentially expressed genes in lesional skin. This connection remains underpowered due to the limited number of subjects with available microbiome data, so results are not shown. *SOD2* (superoxidase dismutase 2) was highly significant in our results and in previous studies of acne (Trivedi *et al*., 2006). *SOD2* codes for an enzyme that modulates skin health by regulating cell proliferation, epidermal thickness, and inflammation (Scheurmann *et al*., 2014; Velarde *et al*., 2012; Weyemi *et al*., 2012). Recent work has shown that *SOD3*, another member of this superfamily, suppressed *C*. acnes-induced inflammation by inhibiting *TLR2* and *NLRP3* inflammasome activation (Nguyen *et al*., 2018). The authors, however, did not examine the influence of *SOD2* in their study. Our data suggests possible correlation of *SOD2* expression and increased inflammation, potentially stemming from increased *C. acnes* burden. Although these findings are preliminary, they warrant further consideration.

Although we conducted comprehensive molecular profiling in acne patients and controls, it is worth pointing out three limitations of this study. First, the study lacked patients with severe acne. It is possible that severe acne patients exhibit systemic immune activation, which was not detected in our mild-to-moderate acne population. Genome-wide association studies have identified risk loci in severe acne patients (He *et al*., 2014; Navarini *et al*., 2014), and future studies should evaluate whether in patients with severe acne there is a link between skin disease and systemic inflammation. Second, we collected skin biopsies for transcriptional and microbiome profiling from two different anatomical locations, the back and face, respectively. Although we observed correlations between host response and microbial abundance, it is likely bacterial diversity on the back differs from the pores of the face. Third, we characterized the follicular microbiome using long-read SMRT DNA sequencing and found a significant association between *C. acnes* abundance and acne status. Although other work has shown increased *C. acnes* abundance in acne patients compared to controls (Jahns *et al*., 2012) consistent with our findings, another study that examined the facial microbiome from swabs and deep, short-read sequences reported differences in ribotypes of *C. acnes* in cases rather than differences in abundance (Fitz-Gibbon 2013). Our samples lacked sufficient sequence coverage to determine ribotypes and explore this hypothesis.

This study reflects the most extensive study of acne patients and non-acne controls. The molecular networks identified from our datasets generated new hypotheses and highlighted new molecular drivers for acne pathogenesis, some of which were validated in this study. Collectively, these data and the accompanying results offer a framework for examining new therapeutic targets and provide a repository of molecular and cellular data to guide future studies.

## MATERIALS AND METHODS

### Ethics statement

This human research study (IF1719992) was approved by the Program for the Protection of Human Subjects of Mount Sinai Medical School of Medicine’s institutional review board (IRB). All individuals provided written informed consent and were able to comply with the requirements of the study protocol.

### Study population and sample collection

We recruited 56 acne patients and 20 non-acne controls (**Table 1**) based on inclusion/exclusion criteria provided in **Text S1**. For each individual a range of sample types were collected (**Figure 1**), including 1) skin tissue biopsies from the back, 2) follicular contents from the face, and 3) whole blood, each described in **Text S2**.

### Molecular assays

We measured RNA expression in skin biopsies via RNA sequencing (RNA-seq) and qRT-PCR. We assayed whole blood via (i) RNA-seq, (ii) DNA genotyping, (iii) protein profiling (Olink), and (iv) CyTOF mass cytometry for cell type profiling. The microbiome was assayed using whole genome sequencing of DNA isolated and amplified from follicular biopsies using Pacific Biosciences SMRT metagenomic sequencing. Detailed methods are described in **Text S3**.

### Analyses

#### Statistical tests and visualization

Unless otherwise noted, we conducted analyses and visualization using the R statistical software v3.4.4 (R Core Team, 2015).

#### Differential gene expression

We tested all genes for significant differential expression using two separate comparisons: (1) between lesion and non-lesion samples in acne patients and (2) between non-lesion samples in acne patients and controls. We adjusted RNA-seq data for possible batch effects. We adjusted for multiple comparisons by controlling the false discovery rate (FDR) using the Benjamini-Hochberg method (Benjamini and Hochberg, 1995), with an FDR of 5%. Up and downregulated genes were determined by ANOVA analyses for all test comparisons.

#### Functional enrichment

We performed gene set enrichment analysis by comparing differentially expressed genes and coexpression modules to gene sets that represent a diverse collection of biological functions and annotations. Briefly, we determined significance by comparing the number of overlapping genes observed to the number expected by chance. We used Fisher’s exact tests to estimate the fold-change and the hypergeometric distribution to calculate p-values. (**Text S4a** for more.)

#### Weighted gene co-expression network analysis (WGCNA)

We constructed molecular networks for three sets of samples—(i) lesion samples from patients, (ii) non-lesion samples from patients, and (iii) non-lesion samples from controls—using the WGCNA package (Langfelder and Horvath, 2008) in R {Langfelder, 2012 #2243}. Briefly, we converted normalized expression data to topological overlap matrices and then to distance matrices, for which genes were clustered into co-expression modules. Hub genes were identified using *softConnectivity* scores in the 95^th^ percentile within each module. (**Text S4b** for more.)

#### DNA

We conducted quality control on the genotype data across all samples using PLINK v1.9 (Chang *et al*., 2015). We tested for group-level differences using logistic regression with a Bonferroni threshold of 5% for significance. (**Text S4c** for more.)

#### CyTOF

Automated cell type discovery (ACDC) (Lee et al., 2017) was performed for 37 surface markers (**Table S6**) to test for differences between patients and controls (**Text S4d**).

#### Protein biomarker

We tested Olink normalized protein expression (NPX) values, which are on a Log2 scale, for differences in abundance levels between patients and controls using the Wilcoxon rank sum test, and then corrected for multiple hypothesis testing by adjusting the p-values using the Benjamini-Hochberg FDR method. (**Text S4e** for more.)

#### Microbiome

To estimate metagenomic abundance, we processed long reads from PacBio, SMRT sequencing data (Pacific Biosciences, Menlo Park, CA) and mapped filtered subreads to the MiniKraken metagenomic database (Oct. 2017, 4 GB version) using the Kraken sequence alignment software (Wood and Salzberg, 2014). We tested for differences in species abundance between acne patients and non-acne controls using the Wilcoxon rank sum test. (**Text S4f** for more.)

## Abbreviations

RNA-seq: RNA sequencing
FDR: False Discovery Rate
*C. acnes*: *Cutibacterium acnes*
FC: fold-change
qRT-PCR: quantitative real-time polymerase chain reaction
LM: lesion module
NLM: non-lesion module.

## ACKNOWLEDGEMENTS

This study was supported by the Acne Cure Alliance and the MCJ Amelior Foundation. The clinical fellows were instrumental for patient intake and sample collection. We thank the Core facilities at Mount Sinai for sample processing and data collection from high-throughput assays.

