## Supplemental Material (Figs, Tables) for "Multiscale analysis of acne connects molecular subnetworks with disease status"

**SUPPLEMENTARY MATERIALS AND METHODS**

### SUPPLEMENTARY FIGURES

#### Supplementary Figure 1. Example photographs of lesional and non-lesional skin sites.

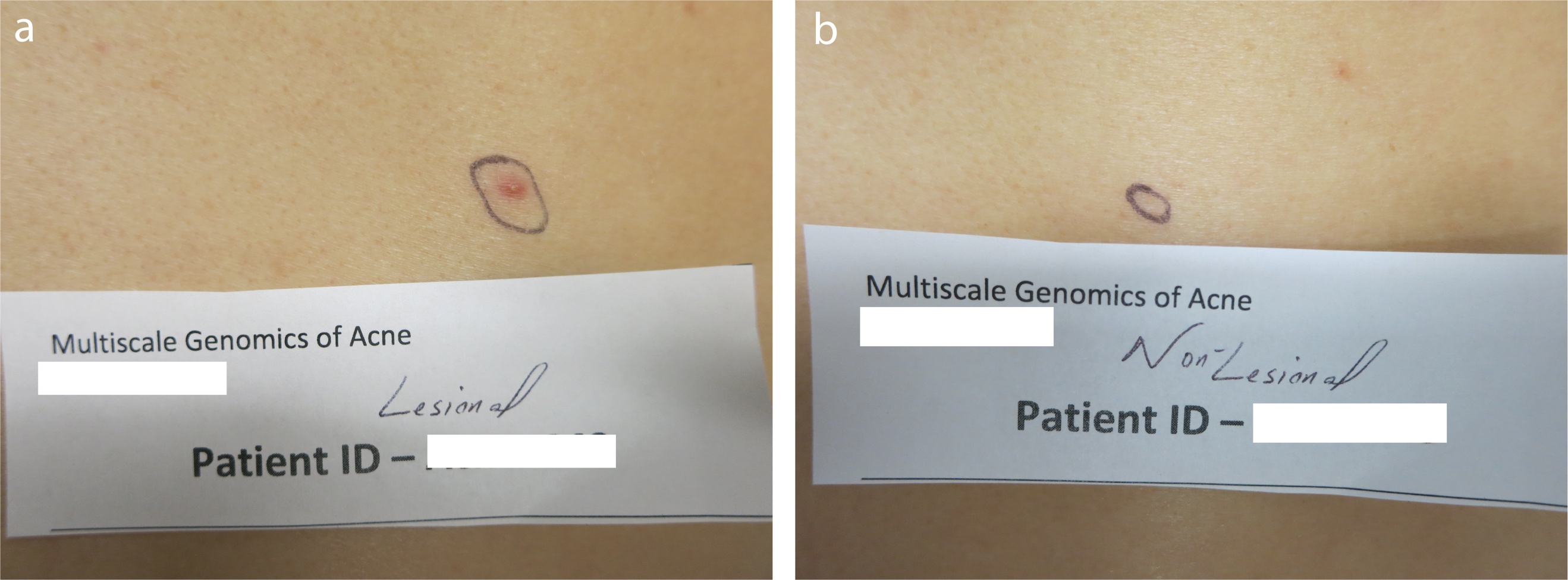

**(a)** Example photographs of lesional and **(b)** non-lesional skin. Biopsy sites were on the upper back in similar areas for all individuals. Non-lesional skin sampling areas were required to be at least 5 mm away from any active acne lesion. Skin biopsy methods are described in **Supplementary Text 2.**

#### Supplementary Figure 2. Manhattan plot of genome-wide single nucleotide polymorphisms.

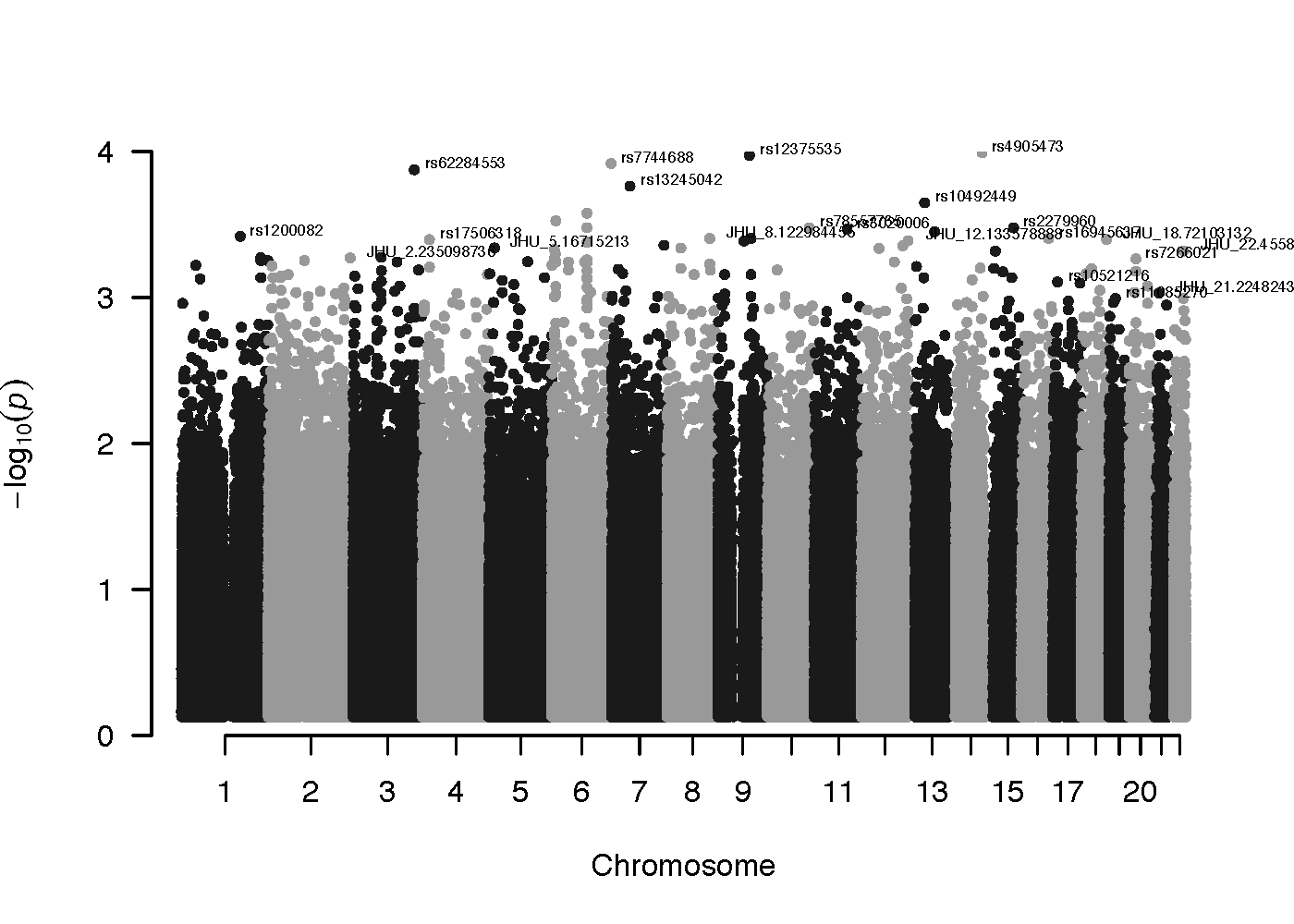

Single Nucleotide Polymorphisms (SNPs) with p > -log_10_(0.0025) are labeled with the SNP name. No SNPs passed the Bonferroni threshold of –log_10_(p) > 7.31, accounting for 1,023,937 autosomal SNPs tested. P-values are from logistic regression, comparing each SNP to association with case status.

#### Supplementary Figure 3. Frequency of immune cell populations in PBMCs determined by CyTOF.

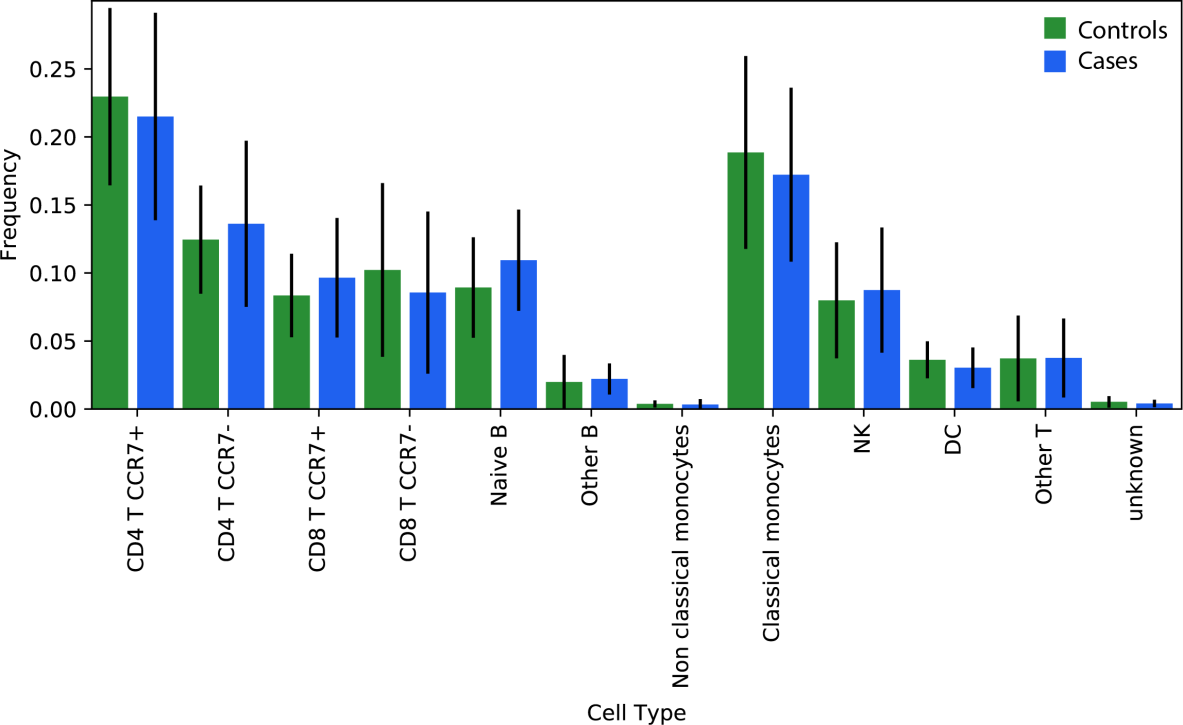

Automated cell type discovery and classification (ACDC). Bars represent the mean for each group; vertical lines represent 1 standard deviation. There were no statistical differences between cases and controls for any of the cell subtypes.

#### Supplementary Figure 4. Genes replicated using qRT-PCR.

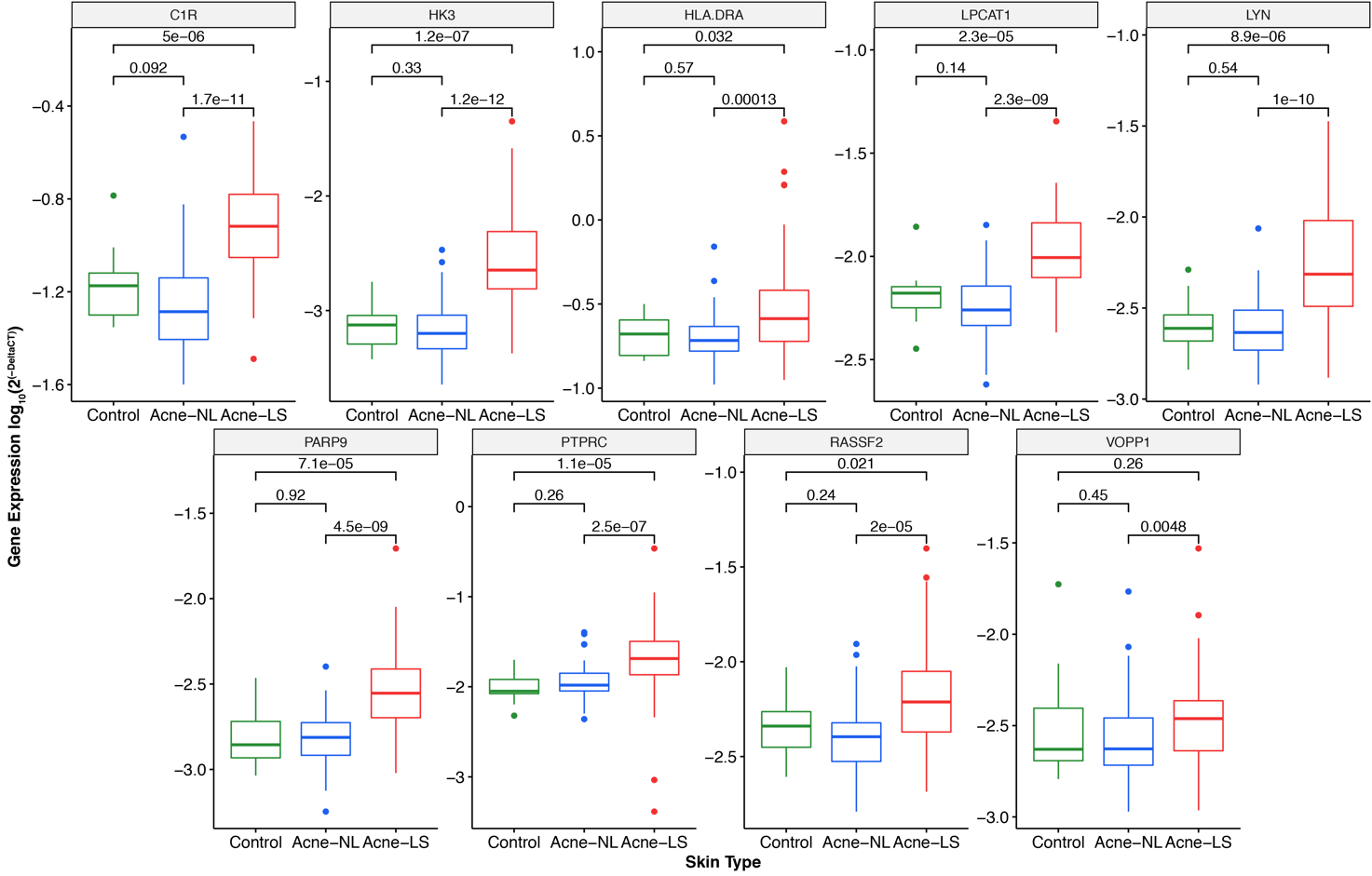

Acne-LS: lesional skin from acne patients; Acne-NL: non-lesional skin from acne patients; Control: non-lesional skin from healthy controls without acne. Unadjusted p-values from Wilcoxon rank sum tests show for each comparison.

#### Supplementary Figure 5. Microbiome sequence read mapping information.

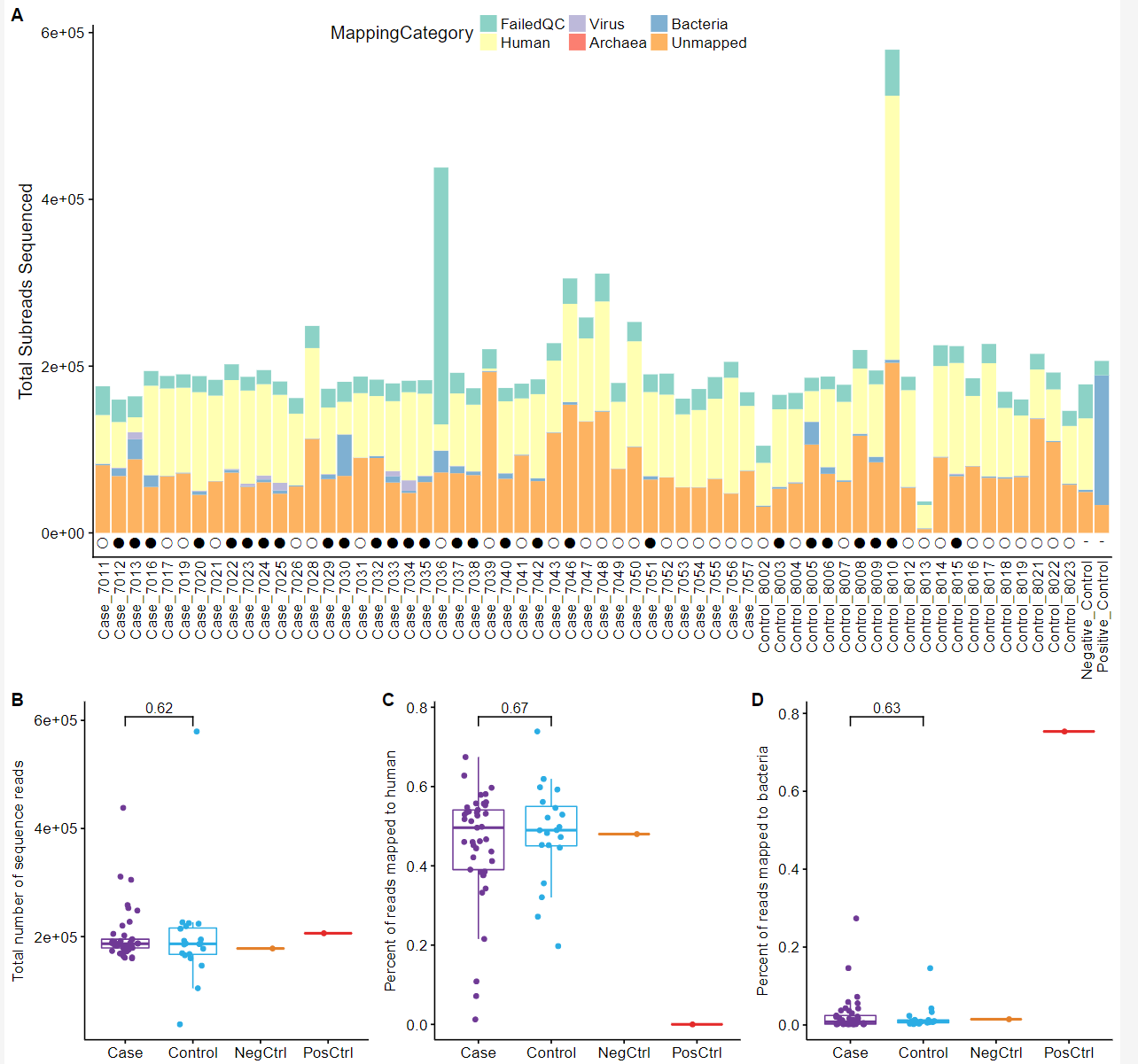

**(a)** Counts for the number of sequence reads that failed quality control, mapped to human/virus/archaea/bacteria, or remained unmapped. Subreads failed QC if they had a length less than 750 bp and/or a quality score less than 0.75. Human reads were mapped and removed from subsequent analyses using BMTagger (Rotmistrovsky and Agarwala 2011). Remaining subreads were mapped to a metagenomic database using Kraken (Wood and Salzberg 2014). Open circles represent samples that failed QC and were not included in analysis. Filled-in circles represent samples that passed QC. **(b)** Total number of sequenced subreads by case status and control type. **(c)** Percent of total reads mapped to human reference. **(d)** Percent of total reads mapped to bacterial genomes. **(b-d)** Each box represents the interquartile range (IQR), horizontal lines in each box represent the median, and the length of the vertical lines (whiskers) represents 1.5 × the upper and lower IQR boundary. Cases: n=41; Controls: n=20; Positive Control: n=1; Negative Control: n=1.

#### Supplementary Figure 6. Species observed in mock community control.

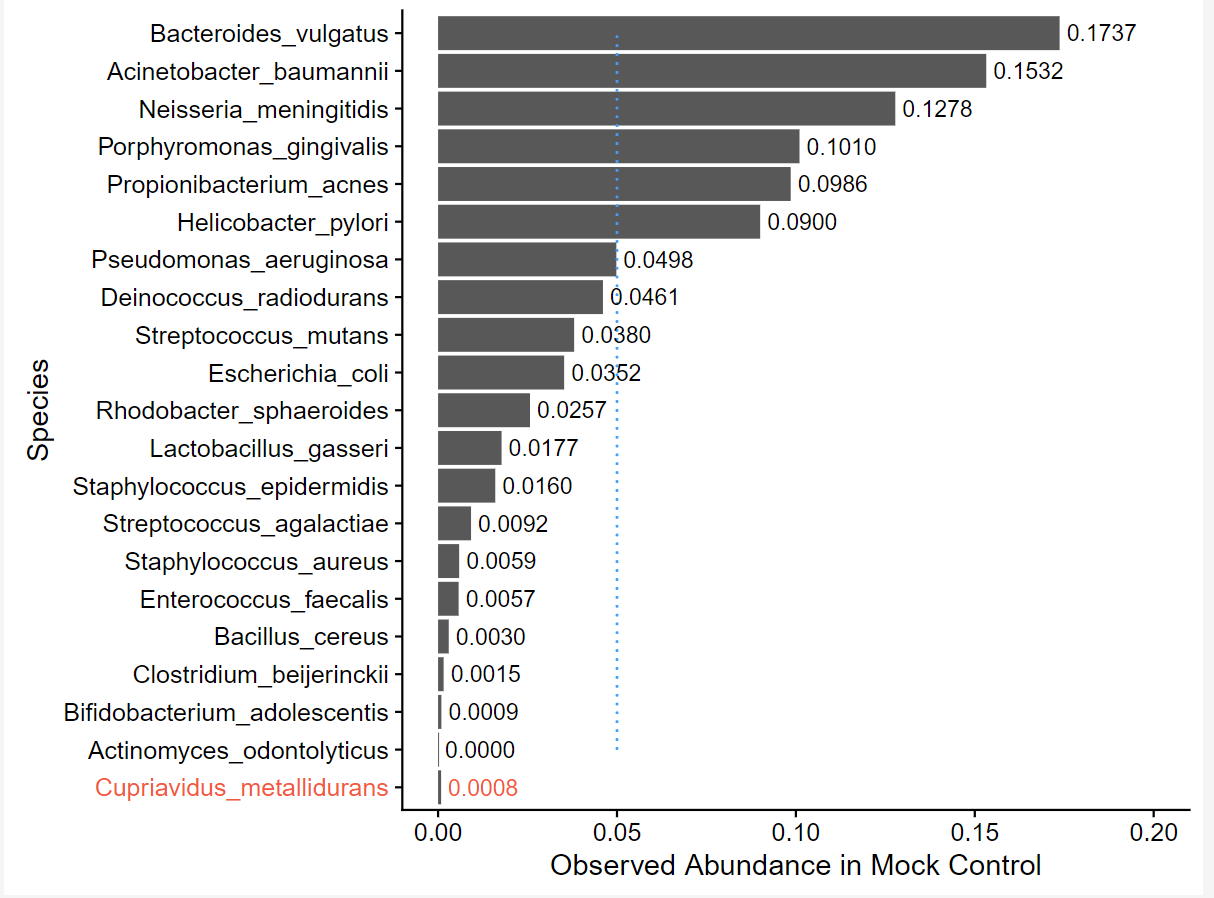

Nineteen of twenty expected species were observed. One species (*Cupriavidus metallidurans*), not expected, was observed at a very low relative abundance (0.08%). The dashed blue line represents the expected abundance (5%) based on the mock control composition (ATCC® MSA-1002™). Species abundance values shown are QC-adjusted.

#### Supplementary Figure 7. Species identification accuracy in mock control.

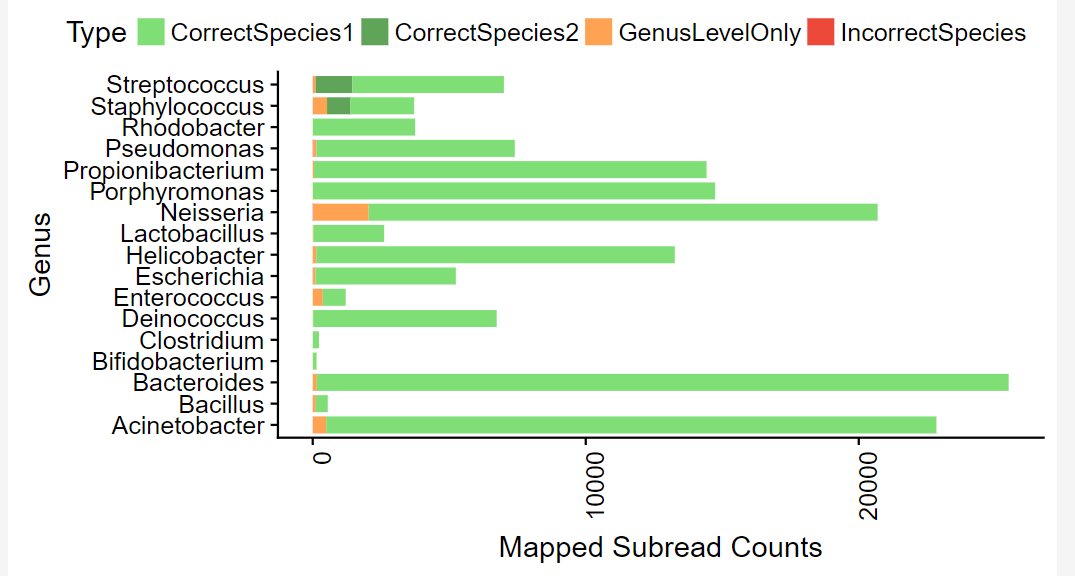

We summarized the ability to speciate correctly based on the sequencing of a mock community (positive) control. Seventeen of 18 genera from the mock community control sample are shown. Actinomyces (1 of 18) is not shown because it was not detected. Genus Cupriavidus (*Cupriavidus metallidurans*, 123 total counts, 0.08% abundance) not shown. The mock community control had two species within each of genus Streptococcus (*S. mutans* – light green; *S. agalactiae* – dark green) and genus Staphylococcus (*S. epidermidis* – light green; *S. aureus* – dark green).

### SUPPLEMENTARY TABLES

#### Supplementary Table 1. Cohort characteristics.

| **Subject ID** | **Case Status** | **Age** | **Sex** | **Race** | **Fitzpatrick Skin Type** | **IGA - Acne Severity Score** |
| --- | --- | --- | --- | --- | --- | --- |
| 7001 | Case | 29 | Female | Caucasian - Non-Hispanic | Type III | 3 - Moderate |
| 7002 | Case | 25 | Female | Hispanic | Type III | 3 - Moderate |
| 7003 | Case | 18 | Female | Caucasian - Non-Hispanic | Type II | 2 - Mild |
| 7004 | Case | 18 | Male | Caucasian - Non-Hispanic | Type III | 1 - Almost Clear |
| 7005 | Case | 21 | Male | Hispanic | Type II | 2 - Mild |
| 7006 | Case | 20 | Male | Hispanic | Type II | 2 - Mild |
| 7007 | Case | 30 | Female | Asian | Type V | 1 - Almost Clear |
| 7008 | Case | 30 | Male | African American | Type VI | 1 - Almost Clear |
| 7009 | Case | 26 | Female | Hispanic | Type III | 2 - Mild |
| 7010 | Case | 19 | Male | Hispanic | Type III | 4 - Severe |
| 7011 | Case | 29 | Female | Hispanic | Type III | 3 - Moderate |
| 7012 | Case | 31 | Male | Hispanic | Type IV | 2 - Mild |
| 7013 | Case | 30 | Male | Hispanic | Type III | 2 - Mild |
| 7014 | Case | 29 | Male | African American | Type VI | 1 - Almost Clear |
| 7016 | Case | 18 | Female | Hispanic | Type III | 2 - Mild |
| 7017 | Case | 31 | Female | Hispanic | Type III | 1 - Almost Clear |
| 7018 | Case | 24 | Female | African American | Type V | 1 - Almost Clear |
| 7019 | Case | 34 | Female | African American | Type V | 2 - Mild |
| 7020 | Case | 32 | Male | Hispanic | Type IV | 2 - Mild |
| 7021 | Case | 31 | Female | Hispanic | Type V | 2 - Mild |
| 7022 | Case | 32 | Female | African American | Type V | 2 - Mild |
| 7023 | Case | 26 | Male | African American | Type V | 3 - Moderate |
| 7024 | Case | 25 | Female | African American | Type V | 2 - Mild |
| 7025 | Case | 27 | Female | African American | Type V | 2 - Mild |
| 7026 | Case | 25 | Male | African American | Type VI | 1 - Almost Clear |
| 7027 | Case | 28 | Male | African American | Type VI | 1 - Almost Clear |
| 7028 | Case | 22 | Male | African American | Type VI | 2 - Mild |
| 7029 | Case | 19 | Male | African American | Type V | 2 - Mild |
| 7030 | Case | 21 | Male | Hispanic | Type III | 3 - Moderate |
| 7031 | Case | 29 | Female | Hispanic | Type III | 1 - Almost Clear |
| 7032 | Case | 22 | Male | African American | Type V | 1 - Almost Clear |
| 7033 | Case | 22 | Male | African American | Type V | 2 - Mild |
| 7034 | Case | 25 | Male | African American | Type V | 3 - Moderate |
| 7035 | Case | 20 | Female | Hispanic | Type IV | 3 - Moderate |
| 7036 | Case | 23 | Male | Hispanic | Type III | 3 - Moderate |
| 7037 | Case | 26 | Male | African American | Type III | 3 - Moderate |
| 7038 | Case | 24 | Male | Hispanic | Type III | 3 - Moderate |
| 7039 | Case | 19 | Male | African American | Type V | 2 - Mild |
| 7040 | Case | 25 | Male | African American | Type V | 3 - Moderate |
| 7041 | Case | 30 | Female | Hispanic | Type II | 3 - Moderate |
| 7042 | Case | 23 | Female | Hispanic | Type IV | 3 - Moderate |
| 7043 | Case | 22 | Female | Hispanic | Type III | 3 - Moderate |
| 7044 | Case | 20 | Male | Mixed | Type III | 2 - Mild |
| 7045 | Case | 21 | Male | Caucasian - Non-Hispanic | Type II | 2 - Mild |
| 7046 | Case | 30 | Female | Asian | Type III | 2 - Mild |
| 7047 | Case | 30 | Male | Caucasian - Non-Hispanic | Type II | 2 - Mild |
| 7048 | Case | 23 | Male | Asian | Type V | 2 - Mild |
| 7049 | Case | 30 | Male | Caucasian - Non-Hispanic | Type II | 2 - Mild |
| 7050 | Case | 22 | Male | Asian | Type IV | 2 - Mild |
| 7051 | Case | 22 | Male | Asian | Type IV | 3 - Moderate |
| 7052 | Case | 24 | Female | Hispanic | Type III | 4 - Severe |
| 7053 | Case | 22 | Male | Hispanic | Type IV | 2 - Mild |
| 7054 | Case | 25 | Female | Hispanic | Type IV | 3 - Moderate |
| 7055 | Case | 25 | Female | Hispanic | Type IV | 2 - Mild |
| 7056 | Case | 22 | Male | African American | Type VI | 2 - Mild |
| 7057 | Case | 25 | Female | Latino | Type III | 3 - Moderate |
| 8002 | Control | 29 | Female | Hispanic | Type III | 0 - None |
| 8003 | Control | 29 | Male | Hispanic | Type IV | 0 - None |
| 8004 | Control | 26 | Male | Caucasian - Non-Hispanic | Type II | 0 - None |
| 8005 | Control | 26 | Male | Caucasian - Non-Hispanic | Type II | 0 - None |
| 8006 | Control | 26 | Male | Caucasian - Non-Hispanic | Type II | 0 - None |
| 8007 | Control | 31 | Male | Hispanic | Type II | 0 - None |
| 8008 | Control | 24 | Female | Hispanic | Type IV | 0 - None |
| 8009 | Control | 24 | Female | Caucasian - Non-Hispanic | Type II | 0 - None |
| 8010 | Control | 28 | Male | Caucasian - Non-Hispanic | Type III | 0 - None |
| 8012 | Control | 35 | Female | Caucasian - Non-Hispanic | Type II | 0 - None |
| 8013 | Control | 23 | Female | African American | Type VI | 0 - None |
| 8014 | Control | 29 | Female | African American | Type IV | 0 - None |
| 8015 | Control | 25 | Female | Hispanic | Type II | 0 - None |
| 8016 | Control | 27 | Female | African American | Type V | 0 - None |
| 8017 | Control | 20 | Female | African American | Type VI | 0 - None |
| 8018 | Control | 26 | Female | African American | Type V | 0 - None |
| 8019 | Control | 34 | Female | Caucasian - Non-Hispanic | Type IV | 0 - None |
| 8021 | Control | 24 | Female | African American | Type V | 0 - None |
| 8022 | Control | 18 | Female | Asian | Type IV | 0 - None |
| 8023 | Control | 34 | Male | Caucasian - Non-Hispanic | Type III | 0 - None |

#### Supplementary Table 2. Proteins assayed using Olink.

| **Protein** | **p*** |  | **Protein** | **p*** |  | **Protein** | **p*** |
| --- | --- | --- | --- | --- | --- | --- | --- |
| 4E-BP1 | 0.1376 |  | FGF-21 | 0.6655 |  | LIF | 0.7195 |
| ADA | 0.3815 |  | FGF-23 | 0.1249 |  | LIF-R | 0.1209 |
| ARTN | 0.2936 |  | FGF-5 | 0.4372 |  | MCP-1 | 0.2697 |
| AXIN1 | 0.0874 |  | Flt3L | 0.8116 |  | MCP-2 | 0.3703 |
| BDNF | 0.1013 |  | GDNF | 0.5387 |  | MCP-3 | 0.0417 |
| Beta-NGF | 0.6578 |  | HGF | 0.3781 |  | MCP-4 | 0.6602 |
| CASP-8 | 0.4363 |  | IFN-gamma | 0.1972 |  | MIP-1 alpha | 0.6707 |
| CCL11 | 0.3710 |  | IL-1 alpha | 0.5348 |  | MMP-1 | 0.3162 |
| CCL19 | 0.6756 |  | IL-10 | 0.9712 |  | MMP-10 | 0.7401 |
| CCL20 | 0.5493 |  | IL-10RA | 0.3279 |  | NRTN | 0.3951 |
| CCL23 | 0.4358 |  | IL-10RB | 0.8566 |  | NT-3 | 0.9137 |
| CCL25 | 0.9424 |  | IL-12B | 0.6237 |  | OPG | 0.4484 |
| CCL28 | 0.4360 |  | IL-13 | 0.8575 |  | OSM | 0.0413 |
| CCL4 | 0.0958 |  | IL-15RA | 0.2334 |  | PD-L1 | 0.8118 |
| CD244 | 0.8739 |  | IL-17A | 0.0836 |  | SCF | 0.9942 |
| CD40 | 0.9030 |  | IL-17C | 0.0494 |  | SIRT2 | 0.2697 |
| CD5 | 0.1032 |  | IL-18 | 0.0419 |  | SLAMF1 | 0.1163 |
| CD6 | 0.6654 |  | IL-18R1 | 0.0348 |  | ST1A1 | 0.0704 |
| CDCP1 | 0.2443 |  | IL-2 | 0.2684 |  | STAMPB | 0.1520 |
| CSF-1 | 0.3040 |  | IL-20 | 0.0790 |  | TGF-alpha | 0.0745 |
| CST5 | 0.9770 |  | IL-20RA | 0.9356 |  | TNF | 0.6501 |
| CX3CL1 | 0.3481 |  | IL-22 RA1 | 0.4671 |  | TNFB | 0.4823 |
| CXCL1 | 0.8967 |  | IL-24 | 0.2905 |  | TNFRSF9 | 0.8619 |
| CXCL10 | 0.2341 |  | IL-2RB | 0.5833 |  | TNFSF14 | 0.6087 |
| CXCL11 | 0.3234 |  | IL-33 | 0.4973 |  | TRAIL | 0.6698 |
| CXCL5 | 0.9654 |  | IL-4 | 0.8789 |  | TRANCE | 0.6549 |
| CXCL6 | 0.4195 |  | IL-5 | 0.7021 |  | TSLP | 0.1625 |
| CXCL9 | 0.7183 |  | IL-6 | 0.0916 |  | TWEAK | 0.2606 |
| DNER | 0.8796 |  | IL-7 | 0.5590 |  | uPA | 0.5151 |
| EN-RAGE | 0.5443 |  | IL-8 | 0.9597 |  | VEGF-A | 0.5025 |
| FGF-19 | 0.8925 |  | LAP TGF-beta-1 | 0.8680 |  |  |  |

Olink targeted circulating protein biomarkers assayed (n=92) and significance for case-control comparisons. *Unadjusted Wilcoxon test p-values shown. After FDR correction, all p-values were 1. Unadjusted values shown for relative comparisons between proteins.

#### Supplementary Table 3. Overlapping differentially expressed genes.

|  | | **Lesion/Non-lesion** | | **Case/Control** | |
| --- | --- | --- | --- | --- | --- |
| **Gene** | **Ensembl ID** | **logFC** | **adj.p** | **logFC** | **adj.p** |
| ***C5orf46*** | ENSG00000178776 | -1.1301 | 4.28E-04 | -0.6811 | 2.70E-02 |
| ***CMTM4*** | ENSG00000183723 | -0.5102 | 1.35E-04 | 0.3257 | 4.78E-02 |
| ***CNKSR2*** | ENSG00000149970 | -0.7542 | 2.02E-02 | -0.7836 | 2.59E-02 |
| ***CP*** | ENSG00000047457 | 0.6232 | 9.96E-03 | -0.7311 | 1.98E-02 |
| ***EHF*** | ENSG00000135373 | 0.4863 | 3.63E-02 | 0.5019 | 4.28E-02 |
| ***EMILIN2*** | ENSG00000132205 | 1.0811 | 3.82E-06 | -0.5279 | 4.77E-02 |
| ***EMX2OS*** | ENSG00000229847 | -0.4265 | 3.43E-02 | -0.4700 | 4.78E-02 |
| ***FYN*** | ENSG00000010810 | 0.4084 | 6.44E-03 | -0.3386 | 4.28E-02 |
| ***KRT16*** | ENSG00000186832 | 1.5638 | 1.74E-03 | 0.9762 | 1.98E-02 |
| ***KRT6A*** | ENSG00000205420 | 1.4040 | 7.79E-03 | 1.1491 | 2.70E-02 |
| ***KRT6C*** | ENSG00000170465 | 2.3244 | 5.24E-04 | 1.2627 | 4.28E-02 |
| ***LILRB5*** | ENSG00000105609 | 1.1754 | 1.40E-04 | -1.0096 | 1.37E-02 |
| ***LTF*** | ENSG00000012223 | 2.7034 | 1.77E-04 | 1.4851 | 3.91E-02 |
| ***MAMDC2*** | ENSG00000165072 | -0.4871 | 2.40E-02 | -0.4134 | 2.59E-02 |
| ***NPR2*** | ENSG00000159899 | -0.4309 | 7.45E-03 | -0.5622 | 1.34E-02 |
| ***S100A8*** | ENSG00000143546 | 2.9937 | 5.72E-05 | 1.8662 | 1.98E-02 |
| ***S100A9*** | ENSG00000163220 | 3.1195 | 4.40E-05 | 2.0298 | 1.37E-02 |
| ***SCN9A*** | ENSG00000169432 | 0.3712 | 3.45E-02 | -0.4206 | 4.73E-02 |
| ***SGCG*** | ENSG00000102683 | -0.9758 | 4.94E-03 | -0.9525 | 2.70E-02 |
| ***SOD2*** | ENSG00000112096 | 1.7436 | 9.21E-08 | 0.4746 | 2.59E-02 |
| ***TMEM64*** | ENSG00000180694 | -0.3643 | 2.30E-03 | 0.3055 | 4.78E-02 |
| ***TTC38*** | ENSG00000075234 | -0.7453 | 4.03E-04 | 0.5416 | 2.70E-02 |
| ***UST*** | ENSG00000111962 | -0.3353 | 4.68E-02 | -0.4418 | 4.45E-02 |
| ***VMP1*** | ENSG00000062716 | 1.2787 | 2.44E-08 | 0.4069 | 4.37E-02 |
| ***ZC3H12A*** | ENSG00000163874 | 0.8878 | 2.73E-04 | 0.5642 | 1.37E-02 |

Data from RNA-seq. Positive fold change corresponds to upregulation in lesion or case samples, respectively. logFC: log fold change; adj.p: FDR-adjusted p-value.

#### Supplementary Table 4. qPCR validation of select RNA-seq results.

|  |  |  | **Lesion/Non-lesion** | | **Case/Control** | |
| --- | --- | --- | --- | --- | --- | --- |
| **Gene** | **EnsemblID** | **TLDA Assay ID** | **qPCR** | **RNA-seq** | **qPCR** | **RNA-seq** |
|  |  |  | **adj.p** | **adj.p** | **adj.p** | **adj.p** |
| ***C1R*** | ENSG00000159403 | Hs00354278_m1 | 3.87E-10 | 1.04E-09 | 0.2839 | 0.8687 |
| ***HK3*** | ENSG00000160883 | Hs01092850_m1 | 5.41E-11 | 1.38E-09 | 0.5340 | 0.8989 |
| ***HLA-DRA*** | ENSG00000204287 | Hs00219575_m1 | 2.40E-04 | 2.57E-04 | 0.6546 | 0.8619 |
| ***LPCAT1*** | ENSG00000153395 | Hs00227357_m1 | 1.35E-08 | 1.04E-09 | 0.3723 | 0.7000 |
| ***LYN*** | ENSG00000254087 | Hs01015816_m1 | 1.19E-09 | 1.04E-09 | 0.6420 | 0.7282 |
| ***PARP9*** | ENSG00000138496 | Hs00967084_m1 | 2.30E-08 | 1.04E-09 | 0.9416 | 0.7880 |
| ***PTPRC*** | ENSG00000081237 | Hs04189704_m1 | 9.66E-07 | 4.33E-06 | 0.4603 | 0.9786 |
| ***RASSF2*** | ENSG00000101265 | Hs00248129_m1 | 4.43E-05 | 2.16E-05 | 0.4412 | 0.8222 |
| ***S100A8*** | ENSG00000143546 | Hs00374264_g1 | 3.09E-06 | 5.72E-05 | 0.0090 | 0.0198 |
| ***S100A9*** | ENSG00000163220 | Hs00610058_m1 | 2.91E-07 | 4.40E-05 | 0.0090 | 0.0137 |
| ***VOPP1*** | ENSG00000154978 | Hs01033979_m1 | 6.10E-03 | 8.30E-05 | 0.5904 | 0.9478 |

adj.p, adjusted p-value; qPRC, qRT-PCR

#### Supplementary Table 5. Differential species abundance between cases and controls.

| **Species** | **p** | **adj.p** | **Cases** | **Controls** |
| --- | --- | --- | --- | --- |
| *Cutibacterium acnes* | 0.0003 | 0.0203 | 58.90% | 2.42% |
| *Cutibacterium phage* | 0.0645 | 0.3043 | 17.21% | 7.04% |
| *Staphylococcus epidermidis* | 0.2092 | 0.5367 | 6.77% | 29.50% |
| *Corynebacterium kroppenstedtii* | 0.0117 | 0.1492 | 2.67% | 0.00% |
| *Streptococcus sanguinis* | 0.4285 | 0.6121 | 2.64% | 0.00% |
| *Stenotrophomonas maltophilia* | 0.6121 | 0.6121 | 2.28% | 0.00% |
| *Streptococcus gordonii* | 0.1048 | 0.3043 | 1.73% | 10.31% |
| *Finegoldia magna* | 0.1035 | 0.3043 | 1.30% | 0.13% |
| *Lactobacillus sakei* | 0.6121 | 0.6121 | 1.25% | 0.00% |
| *Streptococcus mitis* | 0.0007 | 0.0212 | 0.57% | 10.38% |
| *Mycoplasma hyopneumoniae* | 0.1136 | 0.3046 | 0.56% | 2.05% |
| *Streptococcus pneumoniae* | 0.0177 | 0.1492 | 0.29% | 6.20% |
| *Streptococcus pseudopneumoniae* | 0.0025 | 0.0497 | 0.27% | 5.55% |
| *Lactobacillus buchneri* | 0.6121 | 0.6121 | 0.27% | 0.00% |
| *Streptococcus parasanguinis* | 0.4285 | 0.6121 | 0.24% | 2.84% |
| *Pseudomonas stutzeri* | 0.4285 | 0.6121 | 0.20% | 0.00% |
| *Staphylococcus aureus* | 0.5029 | 0.6121 | 0.19% | 0.06% |
| *Staphylococcus haemolyticus* | 0.6121 | 0.6121 | 0.19% | 0.00% |
| *Lactobacillus plantarum* | 0.6121 | 0.6121 | 0.18% | 0.00% |
| *Leuconostoc carnosum* | 0.0848 | 0.3043 | 0.15% | 10.91% |
| *Streptococcus oligofermentans* | 0.0177 | 0.1492 | 0.15% | 3.11% |
| *Leuconostoc kimchii* | 0.6121 | 0.6121 | 0.15% | 0.00% |
| *Dickeya phage RC-2014* | 0.4285 | 0.6121 | 0.15% | 0.64% |
| *Cutibacterium avidum* | 0.6121 | 0.6121 | 0.14% | 0.00% |
| *Lactobacillus helveticus* | 0.6121 | 0.6121 | 0.12% | 0.00% |
| *Sideroxydans lithotrophicus* | 0.6121 | 0.6121 | 0.11% | 0.00% |
| *Mannheimia haemolytica* | 0.0848 | 0.3043 | 0.11% | 1.18% |
| *Pediococcus claussenii* | 0.6121 | 0.6121 | 0.11% | 0.00% |
| *Streptococcus thermophilus* | 0.6121 | 0.6121 | 0.11% | 0.00% |
| *Lactobacillus fermentum* | 0.6121 | 0.6121 | 0.10% | 0.00% |
| *Bacillus megaterium* | 0.6121 | 0.6121 | 0.09% | 0.00% |
| *Corynebacterium diphtheriae* | 0.4285 | 0.6121 | 0.08% | 0.00% |
| *Lactobacillus brevis* | 0.6121 | 0.6121 | 0.08% | 0.00% |
| *Corynebacterium aurimucosum* | 0.4285 | 0.6121 | 0.08% | 0.00% |
| *Streptococcus intermedius* | 0.6121 | 0.6121 | 0.08% | 0.00% |
| *Enterococcus faecium* | 0.6121 | 0.6121 | 0.06% | 0.00% |
| *Pseudomonas fluorescens* | 0.4285 | 0.6121 | 0.05% | 0.00% |
| *Lactococcus lactis* | 0.6121 | 0.6121 | 0.05% | 0.00% |
| *Rothia mucilaginosa* | 0.6121 | 0.6121 | 0.05% | 0.00% |
| *Anaerococcus prevotii* | 0.4285 | 0.6121 | 0.05% | 0.00% |
| *Peptoclostridium difficile* | 0.6121 | 0.6121 | 0.04% | 0.00% |
| *Campylobacter hominis* | 0.6121 | 0.6121 | 0.03% | 0.00% |
| *Filifactor alocis* | 0.6121 | 0.6121 | 0.03% | 0.00% |
| *Lactobacillus casei* | 0.6121 | 0.6121 | 0.02% | 0.00% |
| *Lactobacillus sanfranciscensis* | 0.6121 | 0.6121 | 0.02% | 0.00% |
| *Pseudomonas putida* | 0.6121 | 0.6121 | 0.02% | 0.00% |
| *Lactobacillus salivarius* | 0.6121 | 0.6121 | 0.02% | 0.00% |
| *Cellvibrio japonicus* | 0.6121 | 0.6121 | 0.02% | 0.00% |
| *Lactobacillus kefiranofaciens* | 0.6121 | 0.6121 | 0.01% | 0.00% |
| *Leuconostoc gasicomitatum* | 0.1083 | 0.3043 | 0.00% | 0.62% |
| *Leuconostoc gelidum* | 0.1083 | 0.3043 | 0.00% | 1.60% |
| *Macrococcus caseolyticus* | 0.1083 | 0.3043 | 0.00% | 0.19% |
| *Porphyromonas gingivalis* | 0.0175 | 0.1492 | 0.00% | 1.67% |
| *Prevotella dentalis* | 0.1083 | 0.3043 | 0.00% | 0.07% |
| *Staphylococcus carnosus* | 0.1083 | 0.3043 | 0.00% | 0.23% |
| *Staphylococcus saprophyticus* | 0.1083 | 0.3043 | 0.00% | 0.07% |
| *Staphylococcus warneri* | 0.1083 | 0.3043 | 0.00% | 1.03% |
| *Streptococcus dysgalactiae* | 0.1083 | 0.3043 | 0.00% | 2.13% |
| *Veillonella parvula* | 0.1083 | 0.3043 | 0.00% | 0.09% |

p, p-value from Wilcoxon rank sum test; adj.p, p-value adjusted using B-H FDR; Mean abundance for cases and controls are shown.

#### Supplementary Table 6. CyTOF Antibody Panels.

| **Isotope** | **Target** | **Clone** | **Source** |
| --- | --- | --- | --- |
| **Antibody Panel** | | | |
| 113 In | CD57 | HCD57 | Biolegend |
| 115 In | CD11c | Bu15 | Biolegend |
| 141 Pr | CLA | HECA-452 | Biolegend |
| 142 Nd | CD19 | HIB19 | Biolegend |
| 143 Nd | CD45RA | HI100 | Biolegend |
| 144 Nd | CD62L | DREG-56 | Biolegend |
| 145 Nd | CD4 | RPA-T4 | Biolegend |
| 146 Nd | CD8a | RPA-T8 | Biolegend |
| 147 Sm | CD49d | 9F10 | Biolegend |
| 148 Nd | CD16 | 3G8 | Biolegend |
| 149 Sm | CD127 | A019D5 | Biolegend |
| 150 Nd | CD1c | L161 | Biolegend |
| 151 Eu | CD123 | 6H6 | Biolegend |
| 152 Sm | CD66b | G10F5 | Biolegend |
| 153 Eu | PD-1 | EH12.2H7 | Biolegend |
| 155 Gd | CD27 | O323 | Biolegend |
| 156 Gd | PD-L1 | 29E.2A3 | Biolegend |
| 158 Gd | BTLA | MIH26 | Biolegend |
| 159 Tb | CD103 | Ber-Act8 | Biolegend |
| 160 Gd | CD14 | M5E2 | Biolegend |
| 161 Dy | CD56 | B159 | BD Biosciences |
| 163 Dy | CCR4 | 205410 | R&D Systems |
| 164 Dy | CCR10 | 314305 | Fluidigm |
| 165 Ho | CCR6 | G034E3 | Biolegend |
| 166 Er | CD25 | M-A251 | Biolegend |
| 167 Er | CCR7 | G043H7 | Biolegend |
| 168 Er | CD3 | UCHT1 | Biolegend |
| 170 Er | CD38 | HB-7 | Biolegend |
| 171 Yb | CD161 | HP-3G10 | Biolegend |
| 173 Yb | CXCR3 | G025H7 | Biolegend |
| 174 Yb | HLADR | L243 | Biolegend |
| 175 Lu | CD29 | TS2/16 | Biolegend |
| 176 Yb | CD26 | BA5b | Biolegend |
| **Barcoding** | | | |
| 89 Y | CD45 | HI30 | Fluidigm |
| 194 Pt | CD45 | HI30 | Fluidigm |
| 195 Pt | CD45 | HI30 | Fluidigm |
| 196 Pt | CD45 | HI30 | Fluidigm |
| 198 Pt | CD45 | HI30 | Fluidigm |
| **Cell identification/viability** | | | |
| 103 Rh | Nucleic acid | - | Fluidigm |
| 191/193 Ir | Nucleic acid | - | Fluidigm |

### SUPPLEMENTARY TEXT

#### Supplementary Text 1. Study participant inclusion and exclusion criteria.

**Acne patients (cases)** were adults 18 to 35 years old at the time of consent who had acne symptoms for at least 30 days prior to providing samples. Exclusion criteria included: 1) subjects not willing or able to provide written consent, 2) subjects with scarring or other healing problems, including history of keloids, 3) subjects on acne medications (including topicals, oral medications, or light or laser therapies including home devices, but excluding taxol or lithium) within two weeks prior to the start of the study, 4) subjects who had used oral antibiotics within 30 days of the study visit, 5) use of isotretinoin within one year prior to the study, 6) use of medications that cause chronic immunosuppression (including, but not limited to, oral or injectable corticosteroids, azathioprine, cyclophosphamide, mycophenolate mofetil, tacrolimus, sirolimus, methotrexate, and the use of biological agents including etanercept, adalimumab, alefacept, efalizumab, and infliximab), 7) known sensitivity to lidocaine or topical anesthetic, and 8) women who were pregnant.

**Non-acne controls** were recruited as adults 18 to 35 years old at the time of consent who had no history of acne starting at age 18. Additional exclusion criteria for controls included: 1) scarring or other healing problems, including history of keloids, 2) use of medications that cause chronic immunosuppression (including, but not limited to, oral or injectable corticosteroids, azathioprine, cyclophosphamide, mycophenolate, mofetil, tacrolimus, sirolimus, methotrexate, and the use of biological agents including etanercept, adalimumab, alefacept, efalizumab, and infliximab), 3) known sensitivity to lidocaine or topical anesthetic, and 4) women who were pregnant.

#### Supplementary Text 2. Sample collection procedures.

**Skin biopsies** were performed by trained personnel using standard hygienic procedures; pain was minimized using local anesthesia prior to the procedure. For acne patients, two lesional 4 mm punch biopsies were taken directly from the most involved chronic acne site on the back, spaced between one and four cm apart; two 4 mm non-lesional biopsies were taken at the most normal appearing skin sites at least 5 cm away from any lesion and spaced between one and four cm apart. For healthy controls, the same procedure was performed, but only for two non-lesional sites.

**Whole blood** was drawn from each individual by a trained phlebotomist using standard hygienic procedures. Pain was minimized by using a small-bore needle.

**Microbiome** profiles were surveyed using follicular samples from the face (cheek) of each acne patient and healthy control. Individuals were first instructed to remove any makeup or lotion with a paper towel and water. One drop (15 uL) of cyanoacrylate glue was placed on a sterile glass slide, spread around with a clean pipette tip, pressed against the cheek for 30-45 seconds, and then gently pulled away from the skin. The slide was then placed into a sterile 50 mL conical tube. On the same day, under a dissecting microscope, follicular plugs were removed from each glass slide using sterile microforceps and placed into sterile 1.5 mL microfuge tubes containing 300 uL yeast lysis buffer (MasterPure yeast DNA purification kit; Lucigen, Middleton, WI, USA) and stored at -80°C. Additionally, we used a mock community control (MCC) as a positive control (20 Strain Even Mix Genomic Material, MSA-1002, ATCC, Manassas, VA, USA). Additionally, a negative control was prepared using the same sample preparation protocol, except without applying the glass slide with glue to any patients.

#### Supplementary Text 3. Molecular assay procedures.

**RNA processing for PBMCs and skin biopsies**

**RNA extraction and isolation.** RNA was isolated from peripheral blood mononuclear cells (PBMCs) and skin punch biopsies. For blood cells, we started with an average of 4 million cells per sample to extract RNA from PBMCs. Briefly, we used a Qiagen RNeasy Mini kit (Valencia, CA) and followed the manufacturer’s recommended protocol to isolate the total cellular RNA. A Thermo Scientific NanoDrop 2000 spectrophotometer (Wilmington, DE) was used to determine the purity and quantity of the isolated RNA. We isolated total cellular RNA from skin biopsies by transferring each sample into 700 µl of QIAzol and homogenizing the mixture. After a 5-minute incubation at room temperature, we added 140 µl chloroform and thoroughly mixed the samples. Thawed tissue lysates were centrifuged for 15 minutes at 12,000g at 4 °C. Total RNA was extracted from tissues lysates using the Qiagen miRNeasy Mini Kit (Valencia, CA) following the manufacturer’s protocol. RNA integrity number (RIN) scores were quantified using the Agilent 2100 Bioanalyzer (Wilmington, DE), and found to be 8.79 ± 0.88 and 8.02 ± 1.21 (mean ± SD) for PBMCs and skin tissue respectively. Overall, we isolated 2.89 ± 2.35 µg (mean ± SD) total RNA from PBMCs and 2.34 ± 2.23 µg (mean ± SD) from skin tissue for next generation sequencing.

**RNA library preparation and sequencing.** We prepared RNA sequencing libraries from one microgram of total RNA using the Illumina RNA TruSfeq Kit v2 (San Diego, CA) following the manufacturer’s recommended protocol. Briefly, rRNA was depleted from total RNA using the Invitrogen Ribo-minus kit (Carlsbad, CA) to enrich polyadenylated coding RNA and non-coding RNA. We fragmented the Ribominus RNA in the presence of divalent cations at 94 °C and then converted the fragmented RNA into double-stranded cDNA. After polishing the ends of the cDNA, an adenine base was added at the 3ʹ ends, and we ligated Illumina-supplied specific adapters to the sequences. We applied size selection to the adaptor ligated DNA using AMPure XP beads (New England Biolabs Inc. Ipswich, MA) to get an average size of 250 bp. Following initial size selection, sequences were amplified by PCR for 15 cycles. Amplified DNA was further purified using AMPure XP beads (New England Biolabs Inc. Ipswich, MA) to get the final sequence library. An Agilent 2100 Bioanalyzer was used to determine the insert sizes. A Qubit (Thermo Fisher, Waltham, MA) was used to quantify the DNA concentrations. We layered a pool of 5 barcoded RNA sequence libraries on one of the eight lanes of the Illumina flow cell at appropriate concentrations and then bridge amplified to obtain ≥ 30 million pass filter clusters per sample. Sequencing was conducted using an Illumina HiSeq 4000 platform (San Diego, CA), producing 100 bp paired-end reads.

**RNA sequence mapping.** Raw sequence data was processed and analyzed using TopHat and Cufflinks pipelines (Trapnell et al. 2010; Trapnell et al. 2009). RNA-sequencing reads were aligned to GRCh37 with STAR v2.4.0g1 (Dobin et al. 2013). Reads that uniquely mapped to genes were counted with featureCounts v1.4.4 (Liao et al. 2014) using annotations from ENSEMBL v70 (Flicek et al. 2014). All analysis used log2 counts per million (CPM) following mean-variance stabilization and normalization with voom (Law et al. 2014) as implemented in the limma package (Ritchie et al. 2015). We removed transcripts with fewer than 1 count per 100,000 on average across all samples before statistical analysis. For all comparative analyses and gene set enrichment, Ensembl Gene IDs were mapped to Entrez GeneIDs using information from the HUGO Gene Nomenclature Committee (<http://www.genenames.org>) March 19, 2016 (Wain et al. 2002).

**Quantitative RT-PCR (qRT-PCR).** RNA was extracted for real time PCR using EZ-PCR Core reagents (Life technologies, Grand Island, NY) from 56 lesional and non-lesional skins on acne patients and 17 controls, as previously described (Noda et al. 2015). Taqman Low Density Array/TLDA cards were used for qRT-PCR to analyze mRNA expressions of a panel of inflammatory markers, including innate immunity, general inflammation, Th1, Th2, and Th17/22 pathways. Expression values were normalized to housekeeping gene RPLP0. The primers and probes used were generated by Primer Express Algorithm (Applied Biosystems, Foster City, CA) as previously described (Tintle et al. 2011).

**Additional whole blood assays**

**DNA genotyping.** Using the Illumina Infinium Multi-ethnic beadchip we genotyped 1,779,819 markers for 56 cases and 20 controls.

**Serum profiling using proximity extension multiplex assay (Olink)**. Serum samples for 56 cases and 20 controls were analyzed for a panel of circulating proteins using Olink multiplex assay (Olink Bioscience, Uppsala, Sweden) according to the manufacturer’s instructions. The panel included 92 proteins associated with human inflammatory conditions. Briefly, an incubation master mix containing pairs of oligonucleotide-labeled antibodies to each protein, is added to the samples and incubated for 16 hours at 4°C. Each protein is targeted with two different epitope-specific antibodies increasing the specificity of the assay. Presence of the target protein in the sample would bring the partner probes in close proximity, allowing the formation of a double strand oligonucleotide PCR target. Next day, extension master mix in the sample would initiate the specific target sequences to be detected and generates amplicons using PCR in 96 well plate. For the detection of the specific protein, Dynamic array integrated fluidic Circuit (IFC) 96x96 chip is primed, loaded with 92 protein specific primers and mixed with sample amplicons including 3 inter-plate controls and 3 negative controls. Real time microfluidic qPCR is performed in Biomark (Fluidigm, San Francisco, CA) for the target protein quantification. Data were analyzed using Real time PCR analysis software via ΔΔCt method and NPX (Normalized Protein Expression) manager. Data is normalized using internal controls in every single sample, IPC and negative controls and correction factor and expressed as Log2 scale which is proportional to the protein concentration. One NPX difference equals to the doubling of the protein concentration.

**Mass cytometry (CyTOF).** Cryopreserved Peripheral Blood Mononuclear Cells (PBMCs) were thawed and incubated for 20 minutes at 37°C with Rh103 nucleic acid intercalator (Fluidigm) as a viability dye. Batches of 10 PBMC samples were then labeled with CD45 antibodies conjugated to distinct metal isotopes using a C(5,2) combinatorial barcoding strategy and pooled prior to antibody staining. Each barcoded batch of 10 samples included PBMCs from patient samples and healthy controls to mitigate the effects of potential confounding batch effects on downstream statistical comparisons. Each barcoded pool was then stained with a cocktail of metal labeled antibodies (**Supplementary Table 6**). The cells were then washed and fixed with freshly diluted 1.6% formaldehyde in Phosphate-Buffered Saline (PBS) containing 0.125 nM of Ir intercalator (Fluidigm). The samples were stored at 4°C in PBS until acquisition. Immediately prior to acquisition, the samples were washed once in PBS, once with water, and then resuspended at a concentration of 1 million cells/ml in water containing a 1:2 dilution of EQ 4 element beads (Fluidigm). The samples were acquired on a CyTOF2 mass cytometer equipped with a SuperSampler fluidic system (Victorian Airships, Alamo, CA, USA) at an event rate of < 500 events/second. After acquisition, the data were randomized and normalized using bead-based normalization using Fluidigm’s data processing software. The barcoded samples were deconvolved using Boolean gating, and residual EQ beads and dead cells were excluded prior to downstream data analysis.

**Microbiome**

**Metagenomic sequencing.** Pacific Biosciences, Inc. (Menlo Park, CA) RSII SMRT sequencing was performed on follicular biopsies that underwent isolation of microbial material and WGA amplification using the Qiagen REPLI-g mini kit using manufacturer’s protocol. In total, 41 acne cases, 20 healthy controls, 1 negative control, and 1 mock positive control were sequenced (**Supplementary Figure 1A**). Amplified material was quantified and characterized using Qubit HS-DNA quantification and ample material was used for SMRTbell library preparation for WGS on the RSII instrument.

SMRTbell library preparation and sequencing was performed according to the manufacturer’s instructions and reflects the P6-C4 sequencing enzyme and chemistry, respectively. After fragment purification, the purified DNA sample was taken into DNA damage and end-repair steps. Briefly, the DNA fragments were repaired using DNA Damage Repair solution (1X DNA Damage Repeat Buffer, 1X NAD+, 1 mM ATP high, 0.1 mM dNTP, and 1X DNA Damage Repeat Mix) with a volume of 21.1 µL and incubated at 37ºC for 20 minutes. DNA ends were repaired next by adding 1X End Repair Mix to the solution, which was incubated at 25ºC for 5 minutes, followed by the second 0.45X Ampure XP purification step. Next, 0.75 µM of Blunt Adapter was added to the DNA, followed by 1X template Prep Buffer, 0.05 mM ATP low and 0.75 U/µL T4 ligase to ligate (final volume of 47.5 µL) the SMRTbell adapters to the DNA fragments. This solution was incubated at 25ºC overnight, followed by a 65ºC 10-minute ligase denaturation step. After ligation, the library was treated with an exonuclease cocktail to remove un-ligated DNA fragments using a solution of 1.81 U/µL Exo III 18 and 0.18 U/µL Exo VII, then incubated at 37ºC for 1 hour. Two additional 0.80X Ampure XP purifications steps were performed to remove < 1000 bp molecular weight DNA and organic contaminant and SMRTbell was size selected using Bluepippin technology for molecules > 5kb.

Primer was then annealed to the size-selected SMRTbell with the full-length libraries (80ºC for 2 minutes 30 seconds, followed by decreasing the temperature by 0.1º/s to 25ºC). The polymerase-template complex was then bound to the P6 enzyme using a ratio of 10:1 polymerase to SMRTbell at 0.5 nM for 4 hours at 30ºC and then held at 4ºC until ready for magbead loading, prior to sequencing. The magnetic bead-loading step was conducted at 4ºC for 60-minutes per manufacturer’s guidelines. The magbead-loaded, polymerase-bound, SMRTbell libraries were placed onto the RSII machine at a sequencing concentration of 100 pM and configured for a 240-minute continuous sequencing run. Sequencing was conducted to achieve maximum subread N50 across 1 SMRTcell.

#### Supplementary Text 4. Additional information on analyses of molecular data.

**(a) Gene set enrichment analysis.** We curated 3,295 gene sets from the following libraries: BioCarta, Gene Ontology terms (biological processes, molecular functions, and cellular components), KEGG, Reactome, Wiki pathways, Hallmark, tissue-specific transcriptional signatures, cell-type specific transcriptional signatures, and differentially expressed genes in a large collection of disease gene sets. We applied functional enrichment analysis based on over-enrichment in the expected number of overlapping gene sets between each query gene set and signatures of interest (differentially expressed gene sets or coexpression modules). Briefly, we computed the observed number of “hits” as the overlapping genes between a query gene set and the set of genes in the signature of interest. We compared this number to the expected number given the total number of genes in the query set (n), the total number of genes in the signature of interest (m), and the total number of genes in the background set (N_total_ = genes in background), to calculate the expected number (Expected = m × n / N_total_). The fold change is the ratio of the observed to expected hits. We calculated p-values from hypergeometric distributions using Fisher’s exact tests. To reduce the testing space, we removed query sets with less than three genes. Enrichment calculations were carried out systematically over all gene set libraries and signatures of interest. We controlled for multiple testing by adjusting p-values using B-H FDR with a significance threshold of 10%.

**(b) Coexpression network analysis.** Weighted gene coexpression networks were constructed using the WGCNA package (Langfelder and Horvath 2008), in R. We generated coexpression networks for cases (lesion and non-lesion, separately) and controls (non-lesion only). The normalized gene expression matrix was converted to a topological overlap matrix using function *TOMsimilarityFromExpr*, choosing a network-specific power, selected with function *pickSoftThreshold* as the minimum power required to reach a scale-free topology with an R^2^ greater than 0.80. Networks were generated as “signed hybrid” type. Topological overlap matrices were converted to distance matrices (1 – TOM), and clustered using an adaptive tree cutting algorithm (*cutreeDynamic*), which clusters genes into coexpression modules based on local dendrogram topology, specifying a minimum module size of 30 genes. Modules were then merged when pairwise dissimilarity was less than 0.2 (*mergeCloseModules*). Module eigengenes were calculated from the adjacency matrix using the WGCNA function, *moduleEigengenes*. To identify highly connected genes, we determined intramodular connectivity and labeled a gene as a hub gene if it had *softConnectivity* scores in the 97.5^th^ percentile within the module. *SoftConnectivity* was calculated using the *softconnectivity* function from the WGCNA package (Zhang and Horvath 2005).

**(c) DNA analysis.** PLINK v1.9 (Chang et al. 2015) was used to remove non-standard chromosomes (chromosomes other than 1-22, X, and Y were excluded), SNPs with minor allele frequency less than 1%, and SNPs with genotyping rates less than 95%. We used liftOver (Hinrichs et al. 2006) to convert SNP position information from human genome build GRCh37 to CRCh38. After quality control (QC), 1,023,937 remained for 56 cases and 20 controls. We then performed both a targeted SNP analysis to test for significance in 27 previously-associated acne SNPs from the GWAS catalog (July 2017), as well as an exhaustive genome-wide assessment in our dataset. For the targeted assessment, we selected SNPs within ± 30kb of the target SNPs, which captured 306 SNPs in our dataset.

**(d) Cell-subset profiling (Mass Cytometry, CyTOF).** Thirty-seven surface markers were measured (BTLA, CCR10, CCR4, CCR6, CCR7, CD103, CD11b, CD123, CD127, CD14, CD16, CD161, CD19, CD1c, CD25, CD26, CD27, CD29, CD3, CD38, CD4, CD45, CD45RA, CD49d, CD56, CD57, CD62L, CD64, CD66b, CD73, CD8, CLA, CXCR3, HLA-DR, IntegrinB7, PD-1, and PD-L1) from PBMCs for 54 acne patients and 18 healthy controls. We discarded two channels — CD45RA, due to strong background noise, and CD73, due to no expression in batch 6, 7, and 8. With the remaining markers, we defined cell populations as described in the table below. Automated cell type discovery and classification (ACDC) (Lee et al. 2017) was used to classify these pre-defined cell types. ACDC was applied separately to each batch to reduce batch effects. Linear modeling (adjusting for sex, batch, race, and age) was then applied to test for differences in abundance of these cell types between cases and controls.

| **Cell Type** | **Classification Parameters** |
| --- | --- |
| CD4 CCR7+ | CD3+/CD4+/CD8-/CCR7+/CD19- |
| CD4 CCR7- | CD3+/CD4+/CD8-/CCR7-/CD19- |
| CD8 CCR7+ | CD3+/CD4-/CD8+/CCR7+/CD19- |
| CD8 CCR7- | CD3+/CD4-/CD8+/CCR7-/CD19- |
| Naive B | CD3-/CD19+/CD27- |
| Other B | CD3-/CD19+/CD27+ |
| NK | CD3-/CD56+/CD14-/CD64-/CD19+ |
| Non-classical monocytes | CD3-CD14+/CD64+/CD19-/CD16+ |
| Classical monocytes | CD3-/CD14+/CD64+/CD19-/CD16- |
| DC | CD3-/CD56-/CD14-/CD64-/CD19-/HLADR+ |

**(e) Protein biomarker analysis (Olink).** Fifty-six cases and twenty control were assayed for 92 biomarkers. We removed samples that were automatically flagged as having potential problems during processing (8 cases and 2 controls), leaving 48 cases and 18 controls for analysis. We used Wilcoxon tests to compare abundance in cases and controls.

**(f) Microbiome analysis.** Raw PacBio subread base files (.bax.h5) were initially processed using bash5tools.py (<https://github.com/PacificBiosciences/pbh5tools>); using minimum subread length 750bp, minimum quality score 0.75) to generate FASTQ files. Contaminant human DNA reads were removed by mapping all subreads to the ChGR38 human reference genome using bmtagger (<ftp://ftp.ncbi.nlm.nih.gov/pub/agarwala/bmtagger>). Kraken (Wood and Salzberg 2014) was used to map all filtered subreads to the MiniKraken (Dec. 2014) metagenomic database. Kraken-mpa-report (part of Kraken package) was used to summarize reads mapping to all observed taxonomy. Metagenomic mapping results were processed and visualized using R at the species level. Samples with very high percentages of failed subreads (> 25%) or unmapped subreads (> 65%) were removed from further analysis, as well as samples with less than 100,000 total reads. Species with less than 50 mapped subreads were excluded as likely noise or minor contamination. Species abundance data for all patient samples were adjusted by subtracting out absolute counts of species that were observed in the negative control. Samples with less than 1,000 total non-human mapped reads remaining were removed from further analysis. For statistical comparisons of abundance between cases and controls, Wilcoxon rank sum tests were used and FDR multiple testing corrections were applied as appropriate. All microbiome results visualizations were created in R using the ggplot2 (Wickham 2016).
